# A *Borrelia burgdorferi* LptD Homolog Facilitates Flipping of Surface Lipoproteins Through the Spirochetal Outer Membrane

**DOI:** 10.1101/2022.12.21.521298

**Authors:** Huan He, Ankita S. Pramanik, Selene K. Swanson, David K. Johnson, Laurence Florens, Wolfram R. Zückert

## Abstract

*Borrelia* spirochetes are unique among diderm bacteria in their lack of lipopolysaccharide (LPS) in the outer membrane (OM) and their abundance of surface-exposed lipoproteins with major roles in transmission, virulence, and pathogenesis. Despite their importance, little is known about how surface lipoproteins are translocated through the periplasm and the OM. In this study, we characterized *Borrelia burgdorferi* BB0838, a distant homolog of the OM LPS assembly protein LptD. Using a CRISPR interference approach, we showed that BB0838 is essential for cell growth. Upon BB0838 knockdown, sentinel surface lipoprotein OspA was retained in the inner leaflet of the OM, as determined by its inaccessibility to *in situ* proteolysis but its presence in OM vesicles. The secretion, insertion and topology of the *B. burgdorferi* OM porin P66 remained unaffected. MudPIT quantitative mass spectrometry analysis of the *B. burgdorferi* membrane-associated proteome further confirmed the selective periplasmic retention of surface lipoproteins under BB0838 knockdown conditions. Alphafold Multimer modeling predicted a *B. burgdorferi* LptB_2_FGCAD complex spanning the periplasm. Together, this indicates that BB0838 facilitates the essential terminal step in a distinctive spirochetal lipoprotein secretion pathway that evolved in parallel to the LPS secretion pathway in gram-negative bacteria. Hence, BB0838/LptD_Bb_ represents an attractive target for novel antimicrobials.

## INTRODUCTION

Lipoproteins are ubiquitous membrane-associated proteins in monoderm (e.g., gram-positive) and diderm (e.g., gram-negative) bacteria. After Sec-mediated export through the cytoplasmic or inner membrane (IM), they are posttranslationally modified in a multistep process, yielding an acylated N-terminal cysteine at the start of a disordered tether peptide that anchors them peripherally in leaflets of the membrane lipid bilayer. In diderm bacteria, lipoproteins destined for the outer membrane (OM), such as the prototypical Brown’s lipoprotein Lpp, are known to be extracted by the Lol pathway and transported through the periplasm to be anchored in the periplasmic leaflet of the OM (Tokuda & Matsuyama, 2004, Okuda & Tokuda, 2011, Zückert, 2014). IM lipoproteins avoid recognition by the Lol machinery through interaction with IM phospholipids (Hara *et al*., 2003).

Long thought to be primarily confined to the diderm periplasm, lipoproteins have emerged as important virulence factors at the bacterial surface, i.e., the pathogen-host interface in various gram-negative genera, being involved in biological functions ranging from nutrient acquisition to immune evasion (reviewed in (Wilson & Bernstein, 2016). Prototypical surface display pathways range from the classical type 2 secretion system (T2SS) first described for *Klebsiella* pullulanase PulA (Pugsley *et al*., 1986, d’Enfert *et al*., 1987, D’Enfert & Pugsley, 1989, Pugsley *et al*., 1990, Sauvonnet & Pugsley, 1996, Francetic & Pugsley, 2005), the *Neisseria* T5SS or “autotransporter” pathway secreting NalP (van Ulsen *et al*., 2003, Roussel-Jazede *et al*., 2010, Roussel-Jazede *et al*., 2013), or the SLAM machinery mediating secretion of a family of *Neisseria* surface lipoproteins (Hooda *et al*., 2016). Another prominent example is *Escherichia coli* RcsF, an OM lipoprotein involved in envelope stress signaling, that has been shown to be partially surface exposed while it is sequestered within the cavity of an outer membrane protein (OMP) (Konovalova *et al*., 2014, Cho *et al*., 2014, Rodriguez-Alonso *et al*., 2020).

*Borrelia* spirochetes, the causative agents of vector-borne relapsing fever and Lyme borreliosis, are diderm bacteria with a rather distinct envelope structure. Unlike the gram-negatives – and even some other spirochetal genera such as *Leptospira* and *Brachyspira* – *Borrelia* are deficient in lipopolysaccharide (LPS) biosynthesis pathways and consequently lack LPS in the surface leaflet of the OM (Takayama *et al*., 1987, Fraser *et al*., 1997, Zückert, 2019). Instead, their bacterial surface is dominated by abundant, immunogenic, and serotype-defining surface lipoproteins that are differentially expressed and play major roles throughout subsequent phases of the vector-host transmission cycle (reviewed in (Coburn *et al*., 2021)). Our recent study, using the genetically tractable Lyme disease spirochete *Borrelia burgdorferi* as a model organism, demonstrated that two-thirds of the *B. burgdorferi* type strain B31 lipoproteome localizes to the surface (Dowdell *et al*., 2017). Tether peptide protease accessibility studies for selected model surface lipoproteins indicated that these surface lipoproteins are indeed anchored in the surface leaflet of the OM (Chen *et al*., 2011).

The molecular mechanisms guiding this diverse cohort of about 90 *B. burgdorferi* lipoproteins to the spirochetal surface have slowly come into focus. Using *B. burgdorferi* monomeric OspA and dimeric OspC and *Borrelia turicatae* Vsp1 as model surface lipoproteins in conjunction with a fluorescent localization reporter, mRFPΔ4 (Schulze *et al*., 2010), we showed earlier that their sorting determinants are encoded within the disordered N-terminal tether peptides; yet, sorting rules established for gram-negative periplasmic lipoproteins did not apply (Schulze & Zückert, 2006, Schulze *et al*., 2010, Kumru *et al*., 2010, Kumru *et al*., 2011). Mutations within tether peptides generally led to the mislocalization of surface lipoproteins to the periplasmic leaflet of the OM (Schulze & Zückert, 2006, Schulze *et al*., 2010, Kumru *et al*., 2010, Kumru *et al*., 2011). However, this mislocalization could be rescued by introducing mutations or conditions that destabilized the fold of OspA or OspA-calmodulin fusions, respectively (Schulze *et al*., 2010, Chen & Zückert, 2011). This indicated that surface lipoproteins crossed the OM in an at least partially unfolded conformation. Fittingly, dimeric lipoproteins were shown to assemble on the bacterial surface (Kumru *et al*., 2011). Together, this indicated the requirement for an OM “lipoprotein flippase” machinery that translocated both the polypeptide through the OM while also facilitating the flipping of the lipid anchor from the periplasmic to the surface leaflet (Zückert, 2019).

In their studies of the *B. burgdorferi* OM proteome, Akins and colleagues showed that depletion of the *B. burgdorferi* beta-barrel assembly machinery (BAM) protein BamA led to a significant growth defect and also reduced the abundance of surface lipoproteins in the OM (Lenhart & Akins, 2010). This indicated that proper localization of OM lipoproteins is dependent on BAM, most likely due to the BAM-mediated assembly of an essential integral OMP serving as the OM lipoprotein flippase. This led us to query the predicted essential *B. burgdorferi* OMPeome for any potential function in surface lipoprotein localization using a single plasmid-based non-toxic CRISPR interference (CRISPRi) system (Murphy *et al*., 2022). Here, we show that depletion of BB0838, a distant *B. burgdorferi* homolog of the gram-negative OM LPS translocase LptD, leads to selective retention of surface lipoproteins on the periplasmic side of the OM, while assembly of a sentinel beta-barrel OMP remains unaffected. This indicates that BB0838/LptD_*Bb*_ functions downstream of BAM in facilitating OM translocation of *B. burgdorferi* surface lipoproteins, thereby expanding the capacity of LptD homologs for lipidated substrates.

## RESULTS

### Identification of BB0838 as a putative *B. burgdorferi* OM lipoprotein LptD homolog

To overcome the biophysical obstacle of moving an amphipathic molecule across a lipid bilayer membrane, an OM lipoprotein flippase must have two separate domains: a hydrophilic transmembrane lumen that accommodates polypeptides of various dimensions, and a hydrophobic cavity that shelters the lipid moiety. To narrow down our list of candidate proteins, we used Phyre2 (Kelley *et al*., 2015) to generate structural models of 41 *B. burgdorferi* proteins that had been bioinformatically predicted to localize to the OM and/or assume a β -barrel structure (Kenedy *et al*., 2016). Among this set were BB0795/BamA_*Bb*_ (Lenhart & Akins, 2010) and BB0838, at the time an OMP of unknown function (Kenedy *et al*., 2016).

We modeled the structure of BB0838 using i-Tasser and AlphaFold 2 (Yang & Zhang, 2015, Jumper & Hassabis, 2022) **(Fig. 1)**. i-Tasser built the top model on the *Shigella flexneri* complex of LptD and LptE (C-score -2.22, T-score 0.45 ± 0.15, RMSD 14.9 ± 3.6 Å on PDB accession number 4Q35) despite less than 20% amino acid identity between them **(Fig. 1A)**. In gram-negative bacteria, LptD together with lipoprotein LptE form the OM translocon for transporting LPS across the OM and displaying it on the cell surface (reviewed in (Konovalova *et al*., 2017)). The LptD N terminus folds into a β -jellyroll domain with 20 antiparallel β -strands forming a hydrophobic groove extending into the periplasm while the LptD C terminus forms a β -barrel transmembrane domain with 26 antiparallel β -strands (Dong *et al*., 2014). The LPS lipid A moiety is directly inserted into the OM via LptD’s N-terminal hydrophobic core while the hydrophilic oligosaccharide core and polysaccharide O antigen moiety of LPS enter the OM through the hydrophilic transmembrane lumen of the LptD β -barrel. LptE is anchored in the periplasmic leaflet of the OM and inserts into the LptD β -barrel lumen to facilitate the re-orientation of LPS to the bacterial surface (Chimalakonda *et al*., 2011, Chng *et al*., 2010, Freinkman *et al*., 2011, Ruiz *et al*., 2010, Wu *et al*., 2006).

**FIG. 1.**
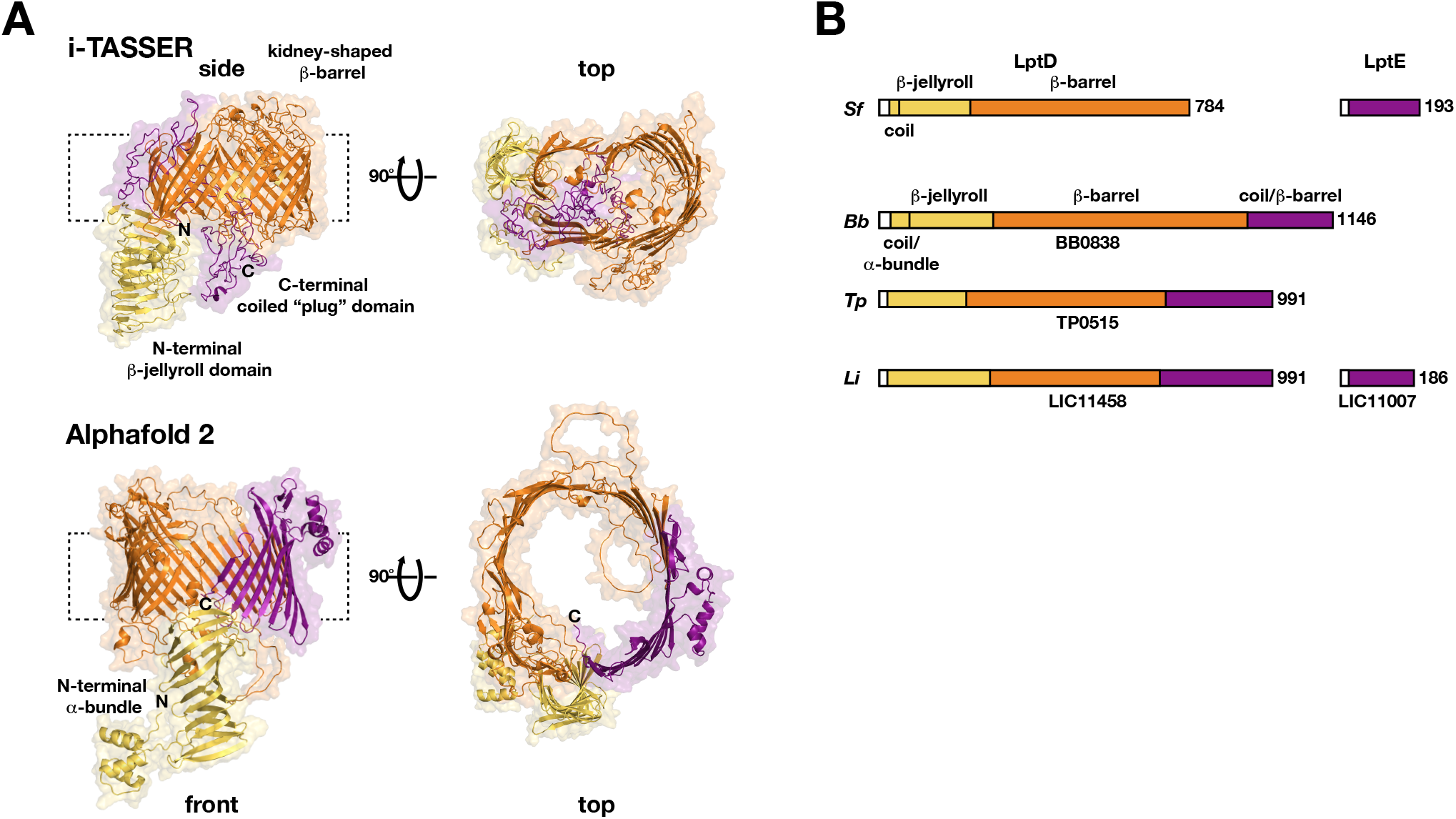
Structural Modeling of BB0838. (A) i-Tasser and AlphaFold models of BB0838. Both algorithms predict a BB0838 fold similar to LptD. In both models, the N-terminal periplasmic β -jellyroll domain is colored in yellow, the central β -barrel domain in orange, and the C-terminal extensions forming either a coiled “plug” or β -barrel extension domains in purple. The i-Tasser model is shown in a side view emphazising the predicted plug domain, and the Alphafold model is shown in a front view emphasizing its separate N-terminal α-bundle and β -jellyroll domains. Both top views are rotated 90° along the vertical axis. The amino and carboxy termini of the protein are indicated by N and C, respectively. Stipled brackets indicate the approximate position of the OM lipid bilayer. **(B)** Comparison of predicted domain structures of spirochetal LptD homologs to gram-negative LptD. Color coding is as in panel A. *Sf, S. flexneri; Bb, B. burgdorferi; Tp, T. pallidum; and Li, L. interrogans*. ORF numbers for each homolog are indicated below each domain bar, and numbers indicate total preprotein length in amino acids. Note that only LPS-containing *L. interrogans* is predicted to have an LptE homolog.

The BB0838 model predicted a slightly shortened N-terminal β -jelly roll domain composed of 18 antiparallel β -strands preceded by a coiled coil. Compared to *S. flexneri* LptD, the β -barrel domain expanded to 28 antiparallel β -strands due to 94 additional amino acids, suggesting an expanded transmembrane lumen that may accommodate larger cargo. In addition to a predicted larger β -barrel, BB0838 has an additional 362 amino acid C-terminal extension that i-Tasser modeled as a coiled coil on LptE within the LptD lumen (Fig. 1). The AlphaFold model of BB0838 **(Fig. 1A, Supplemental Table S3)** similarly predicted a β -jelly roll domain of 18 antiparallel β -strands, but the extreme N terminus assumed an additional domain consisting of a cluster of four short α-helices connected to the β -jelly roll via a flexible linker. In another contrast to the i-Tasser model, BB0838’s C-terminal extension was incorporated into the β -barrel as additional β -strands, leading to a 32-stranded β -barrel that was partially restricted by a large periplasmic loop. In both models, the β -barrel had a lateral opening aligned with the periplasmic β -jelly roll domain. We therefore concluded that BB0838 indeed represents a structural LptD homolog that–in the absence of lipidated polysaccharides within the system–may have evolved to translocate lipidated proteins instead. Intriguingly, even distant LptD homologs of other spirochetal model organisms such as *Treponema pallidum* and *Leptospira interrogans* show a similar domain structure with a C-terminal extension, but only the LPS-containing *L. interrogans* also encodes for an LptE homolog (Hawley *et al*., 2021) (**Fig. 1B, Supplemental Table S1**).

### Generation of a conditional BB0838 knockdown strain using CRISPR interference

A *B. burgdorferi* Tn mutagenesis screen indicated that BB0838 is essential for cell viability (Lin *et al*., 2012, Lin *et al*., 2014). BB0838 is the last gene in a 3-gene operon downstream of *uvrB* and *uvrA* **(Fig. 2A)**, which appear non-essential for growth based on the analysis of a *ΔuvrA* strain (Sambir *et al*., 2011). We therefore generated a conditional *bb0838* knockdown strain using a fully inducible, non-toxic CRISPR interference (CRISPRi) system encoded by a single recombinant *E. coli/B. burgdorferi* shuttle vector, pJJW101 (Murphy *et al*., 2022). pJJW101 carries a *B. burgdorferi* codon-optimized version of dCas9-myc under standard P_QE30_ (T5/*lac* hybrid promoter) control, which avoids toxicity of the original *Streptococcus pyogenes* dCas9 in *B. burgdorferi* under higher induction conditions (Takacs *et al*., 2020). Stringency is further maximized by placing the single guide RNA (sgRNA) module under *trc* (*trp/lac* hybrid) promoter control. Gene-specific target sgRNA sequences were designed with the web-based CRISPy-web interface on an artificial contig assembly of the *B. burgdorferi* B31 chromosome and plasmids to eliminate potential off-target effects as described (Murphy *et al*., 2022, Blin *et al*., 2016). Four target sgRNAs, complementary to either the preferred non-template (NT) strand or to the non-preferred template (T) strand (**Fig. 2B and Table 1**), were then ligated into the pJJW101 sgRNA module. NT1 and NT2 sgRNAs targeting an overlapping sequence close to BB0838’s 5’ end led to a marked *in vitro* growth defect upon CRISPRi induction, whereas T1 and T2 sgRNAs targeting separate sequences further downstream appeared to grow normally (data not shown).

**FIG. 2.**
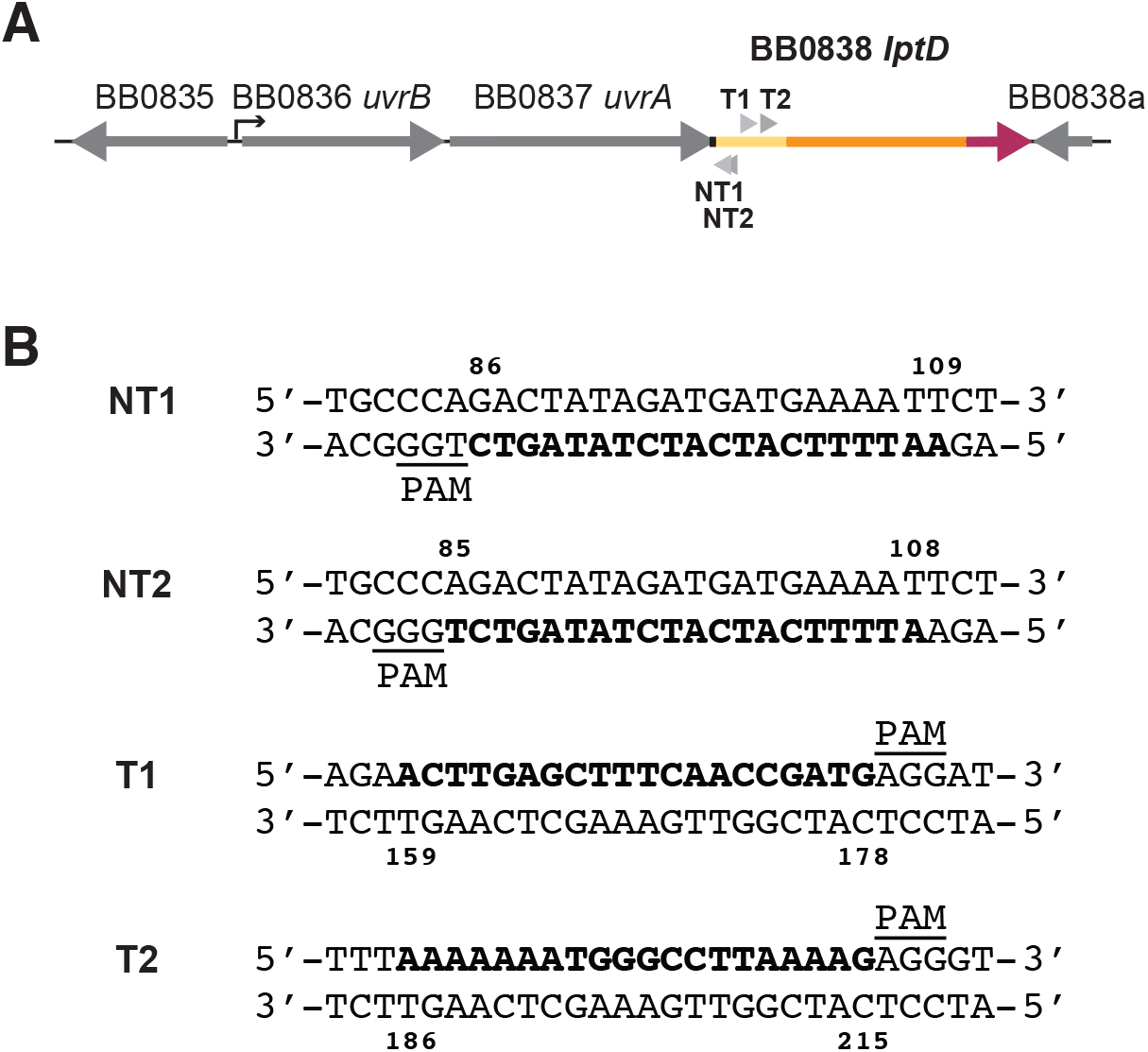
BB0838 operon structure and sgRNAs selected for CRISPRi. **(A)** BB0838 is the last gene in a 3-gene operon with *uvrB* and *uvrA. uvrA* and BB0838 coding sequences overlap by 46 bps, i.e., solely within the signal I leader peptide of BB0838. Arrow heads show the position of all designed sgRNAs within the coding sequence for BB0838’s N-terminal β -jellyroll domain. **(B)** NT1, NT2, T1 and T2 sgRNA sequences are shown in bolded letters, with their start and end nucleotide positions within BB0838 indicated. Associated PAM sequences are underlined.

**Table 1.**
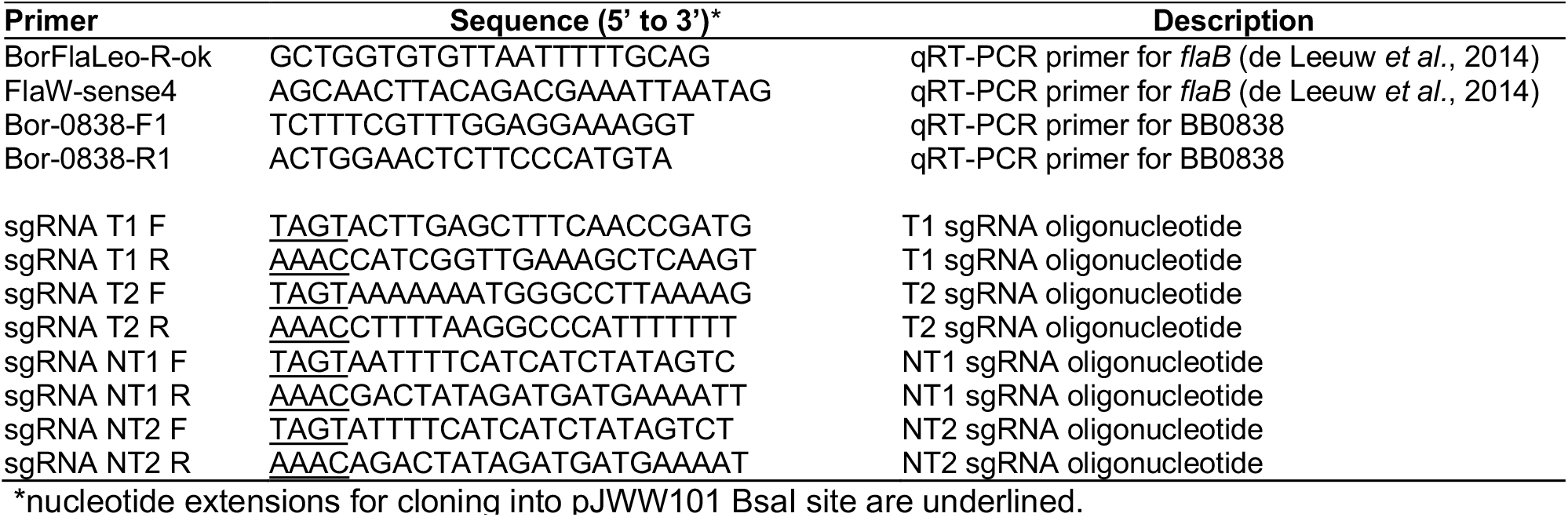
Oligonucleotide primers used in this study.

To evaluate the efficiency of the NT1 and NT2 CRISPRi-mediated BB0838 knockdowns, we used quantitative reverse transcription polymerase chain reaction (qRT-PCR) assays, comparing transcription levels of BB0838 and a *flaB* normalization control in both the mock control strain (transformed with a sgRNA-deficient “empty” pJJW101 plasmid) and in the BB0838 knockdown strains (transformed with the respective sgRNA-containing pJJW101 vectors). IPTG-driven induction of dCas9 and these two sgRNAs led to an 84- and 12-fold knockdown of BB0838 mRNA transcript compared to the mock control, respectively, in total RNA samples collected at day 2 (**Fig. 3**). Due to its higher knockdown efficiency, we selected the NT1 sgRNA-expressing clone as the BB0838 knockdown (KD) strain for further study.

**FIG. 3.**
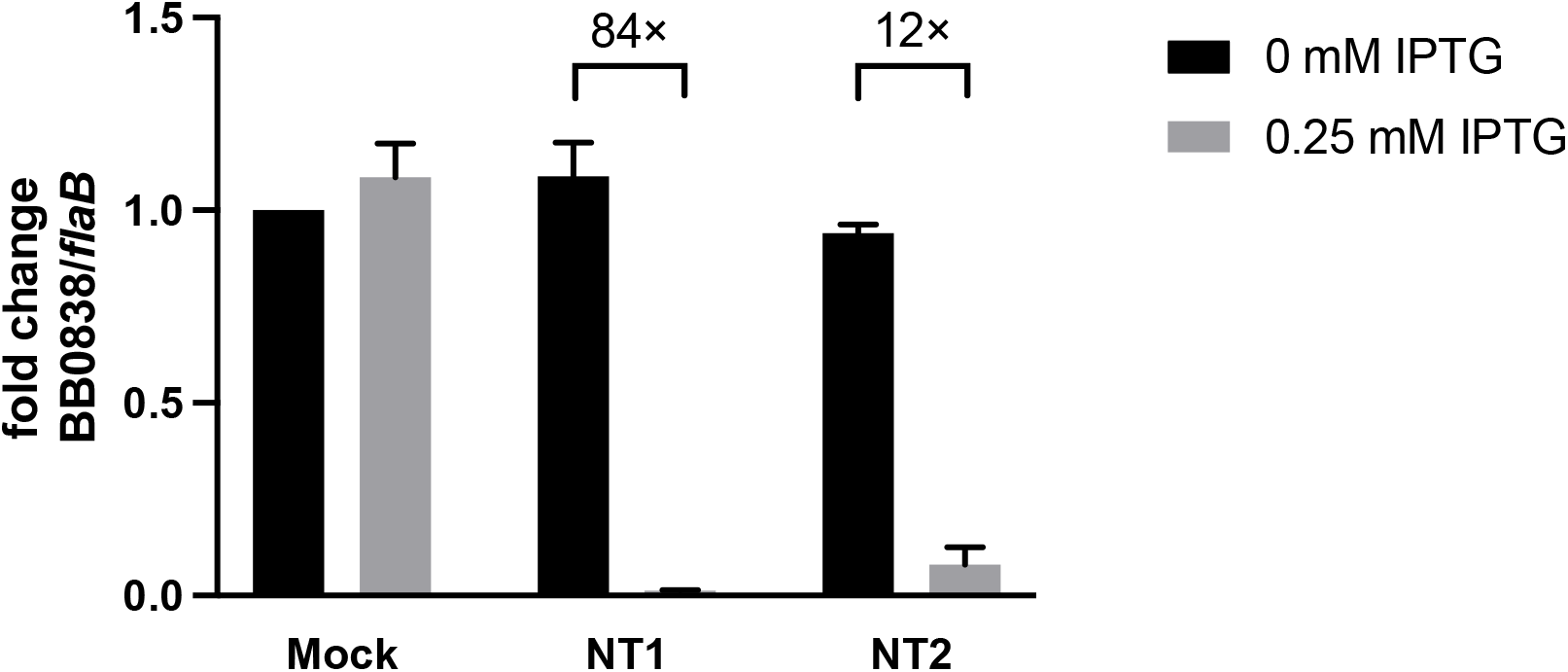
Transcriptional BB0838 knockdown efficiency of NT1 and NT2 sgRNAs. Total RNA samples obtained from *B. burgdorferi* cells harvested at 2 days post-inoculation were assayed by qRT-PCR using specific primers for BB0838 and *flaB*, which was used as a normalization control. Mock, cells harboring “empty” pJJW101 without sgRNA; NT1 and NT2, cells harboring pJJW101 with NT1 or NT2 sgRNAs, respectively. Results are from 3 biological replicates; error bars indicate mean ± SD.

### BB0838 is essential for cell viability

To test for the expected role of BB0838 in cell viability, we monitored cell growth and phenotypic changes of *B. burgdorferi* BB0838 KD cells over time, using non-depleted and wild type cells as a control. 1 × 10^5^ cells/ml were inoculated in complete BSK-II medium with or without 0.25 mM IPTG and followed for 3 days post-inoculation by phase contrast microscopy. As shown by the growth curves in **Fig. 3A**, a significant growth defect in the induced BB0838 KD cultures emerged after day 1, with a 10-fold reduction in cell numbers at day 2 compared to WT that increased to a 100-fold drop by day 3. Higher inducer concentrations did not lead to more severe growth defects (data not shown). Phase contrast micrographs **(Fig. 3B)** showed that depletion of BB0838 led to envelope disturbances and blebbing. The abnormal envelope structure is most likely caused by the envelope stress response resulting from the accumulation of mislocalized surface lipoprotein in the absence of BB0838. Together, these data suggested that dilution of BB0838 to about 10-20% of its WT level is sufficient to disrupt envelope biogenesis and leads to a cessation of bacterial growth.

### BB0838 depletion affects the localization of a major outer surface lipoprotein

To test our hypothesis that BB0838 plays a crucial role in surface lipoprotein translocation, we used proteolytic shaving to assess any changes in surface exposure of lipoproteins in both mock control and BB0838 KD cells. Cultures inoculated with freshly cultured stationary phase cells at 1×10^6^ cells/ml final concentration were grown in selective BSK-II medium with or without 0.25 mM IPTG for 2 days. Harvested and washed intact cells were then treated *in situ* with proteinase K to remove surface-exposed proteins or peptides as described (Bunikis & Barbour, 1999, Zückert *et al*., 2004, Schulze & Zückert, 2006, Chen *et al*., 2011). As shown by Western immunoblotting (**Fig. 4A)** surface lipoprotein OspA was accessible to proteinase K and degraded in both mock and uninduced BB0838 KD cells, while it remained largely protected in the induced BB0838 KD. At the same time, the topology of the OM porin P66, as assessed by protease accessibility of a surface-exposed loop (Bunikis *et al*., 1996, Skare *et al*., 1997, Bunikis & Barbour, 1999, Kenedy *et al*., 2014), remained unaffected. Periplasmic FlaB and cytosolic protein dCas9-Myc served as internal cell integrity controls.

**Fig. 4.**
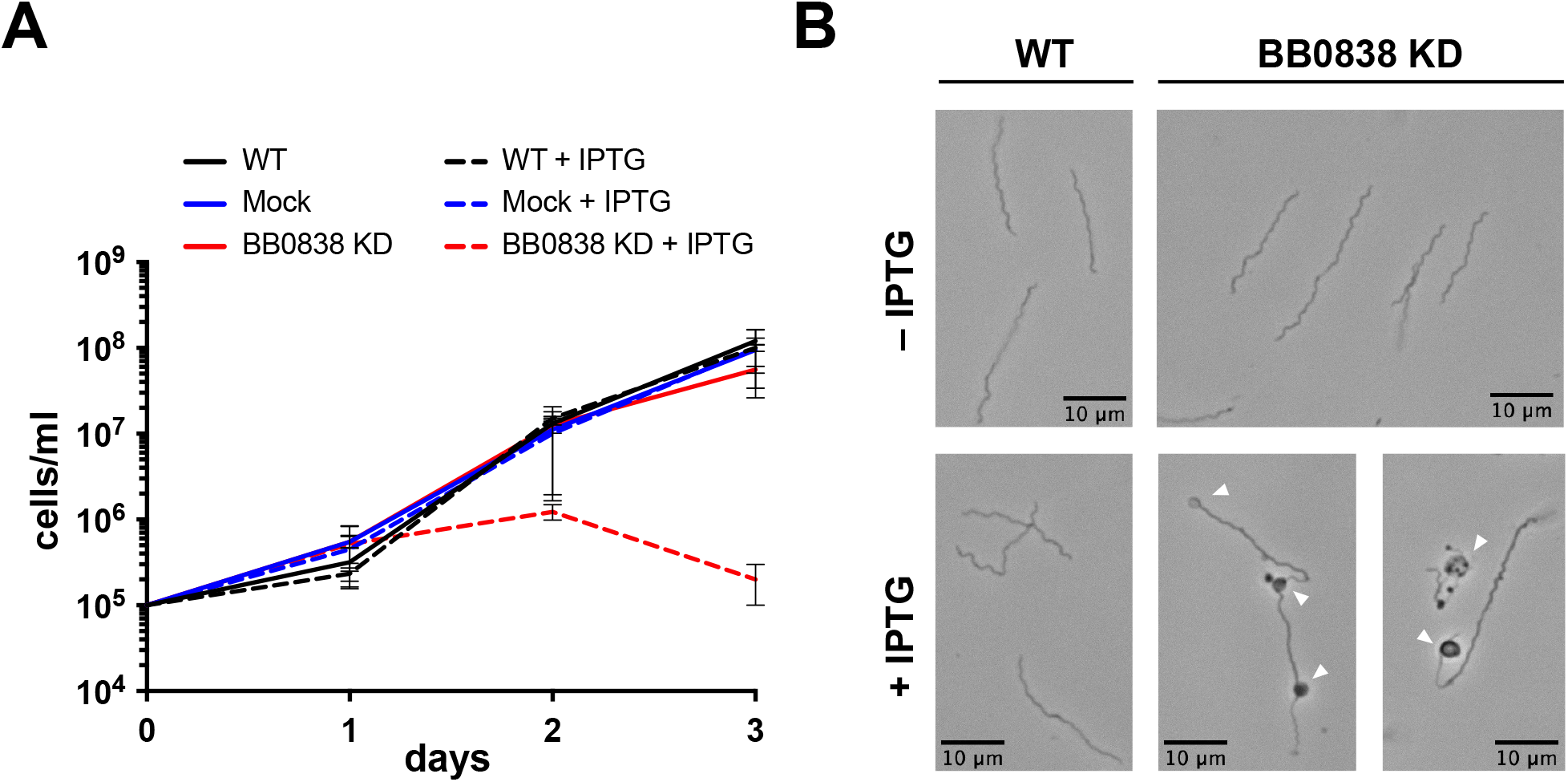
Assessment of BB0838 knockdown-dependent growth defects. **(A)** Growth curve of wild-type (WT), mock control (with “empty” pJJW101) and BB0838 knockdown (KD) cells under inducing (0.25 mM IPTG) and non-inducing conditions. Cells were counted under phase contrast in a Petroff-Hausser counting chamber. Growth curves are from 3 parallel biological replicates; error bars indicate mean ± SD. **(B)** Representative phase micrographs of wild-type (WT) and BB0838 KD *B. burgdorferi* cells at day 2 post-inoculation under both non-inducing and inducing (+0.25 mM IPTG) conditions. White arrowheads indicate visible disturbances in the spirochetal envelope. Size bars are 10 nm.

To assess any defects in surface lipoprotein transport from the IM to the OM under BB0838-depleting conditions, we purified and analyzed the protein content of outer membrane vesicles (OMVs). *B. burgdorferi* cells cultured and harvested as described above at day 2 were osmotically shocked by resuspension in hypotonic citrate buffer, and the released OMVs were purified from remaining cell material (intact cells and protoplasmic cylinders) by ultracentrifugation on a discontinuous sucrose gradient as described (Skare *et al*., 1995, Radolf *et al*., 1995, Dowdell *et al*., 2017). Western immunoblots showed that OspA localized to OMVs under both mock control and BB0838-depleting conditions. OMV purity was assessed by the absence of IM lipoprotein OppAIV (**Fig. 4B**). Together, this set of experiments indicated that depletion of BB0838 primarily blocks the translocation of surface lipoproteins through the OM, but does neither significantly affect their transport through the periplasm, nor disturb the proper topology of integral OMPs such as P66.

### BB0838 depletion leads to a specific localization defect of surface lipoproteins

To evaluate if the observed BB0838-dependent localization defect of OspA extends to the surface lipoproteome in general, we used quantitative multidimensional protein identification technology (MudPIT) mass spectrometry. *B. burgdorferi* mock control and BB0838 KD cells were cultured, harvested, and subjected to proteolytic shaving as described above. Samples were enriched for membrane-associated proteins by Triton X-114 detergent extraction followed by a 2-step precipitation with acetone and trichloroacetic acid as described (Dowdell *et al*., 2017). Two biological replicates were submitted for MudPIT analysis and processed in 5 or 6 technical replicates. We detected 44 of the previously localized 85 lipoproteins encoded by *B. burgdorferi* clone B31-e2 (Dowdell *et al*., 2017). Ten of the undetected lipoproteins (BB0456, BB0475, BB0735, BB0844, BBA33, BBA65, BBB08, BBJ01, BBP39, BBS41) were previously shown to be not transcribed in mid-exponential phase (Arnold *et al*., 2016), and the expression level of the remaining 31 lipoproteins was most likely below the MudPIT detection limit. 13 periplasmic IM, 5 periplasmic OM, and 10 surface lipoproteins were detected in at least two replicates of the uninduced control (838C) and analyzed further (**Supplemental Table S2**).

Based on the normalized spectral abundance factor (dNSAF) as a measure of protein abundance, we calculated the average lipoprotein surface exposure by dividing each protein’s dNSAF in the untreated (–pK) samples by the dNSAF in the corresponding protease-treated (+pK) samples, resulting in a-pK/+pK dNSAF ratio for each protein. Lipoproteins that localize to the periplasm, i.e., are protected from proteinase K, have close to identical dNSAF values in both the -pK and +pK samples and thus dNSAF ratios close to 1, whereas protease-accessible surface lipoproteins show a dNSAF ratio larger than 1 due to lower dNSAF values in the +pK sample (Dowdell *et al*., 2017). Conditional retention of a surface lipoprotein in the periplasm could therefore be followed by a drop of its -pK/+pK dNSAF ratio towards a value of 1.

As expected from our earlier studies, the dNSAF ratios for surface and periplasmic lipoproteins showed a clear separation (**Fig. 6, Supplemental Table S2 and Fig. S1**). dNSAF ratios for all 28 lipoproteins in the control uninduced (838C) samples generally tracked the values obtained with the proteomically more complex *B. burgdorferi* B31 clone B31-A3 (Dowdell *et al*., 2017). The 18 periplasmic IM and OM lipoproteins had dNSAF ratios around 1 (range 0.38 to 2.160. Calculated ratios for surface lipoprotein ranged between 7.12 and 97.63, with three proteins having infinite (∞) ratios due to undetectable peptides in the protease-treated samples, and one known low outlier (BBD10). In the induced BB838 knockdown (838KD) sample, the +pK/-pK dNSAF ratios for the periplasmic IM and OM lipoproteins remained stably around 1 (range 0.20 to 1.83), indicating their continued periplasmic localization. Surface lipoprotein ratios, however, collapsed from their high values in the 838C controls to a range of 0.00 to 3.98, indicating that they were now substantially retained within the periplasm. For comparison to the immunoblot data shown in Fig. 5, OspA showed a drop in +pK/-pK ratio from 45.83 to 1.81. Together, these data indicate that depletion of BB0838 leads to a protein localization defect in *B. burgdorferi* cells that is specific to surface lipoproteins. This implicates BB0838 as the terminal component of a surface lipoprotein secretion pathway.

**Fig. 5.**
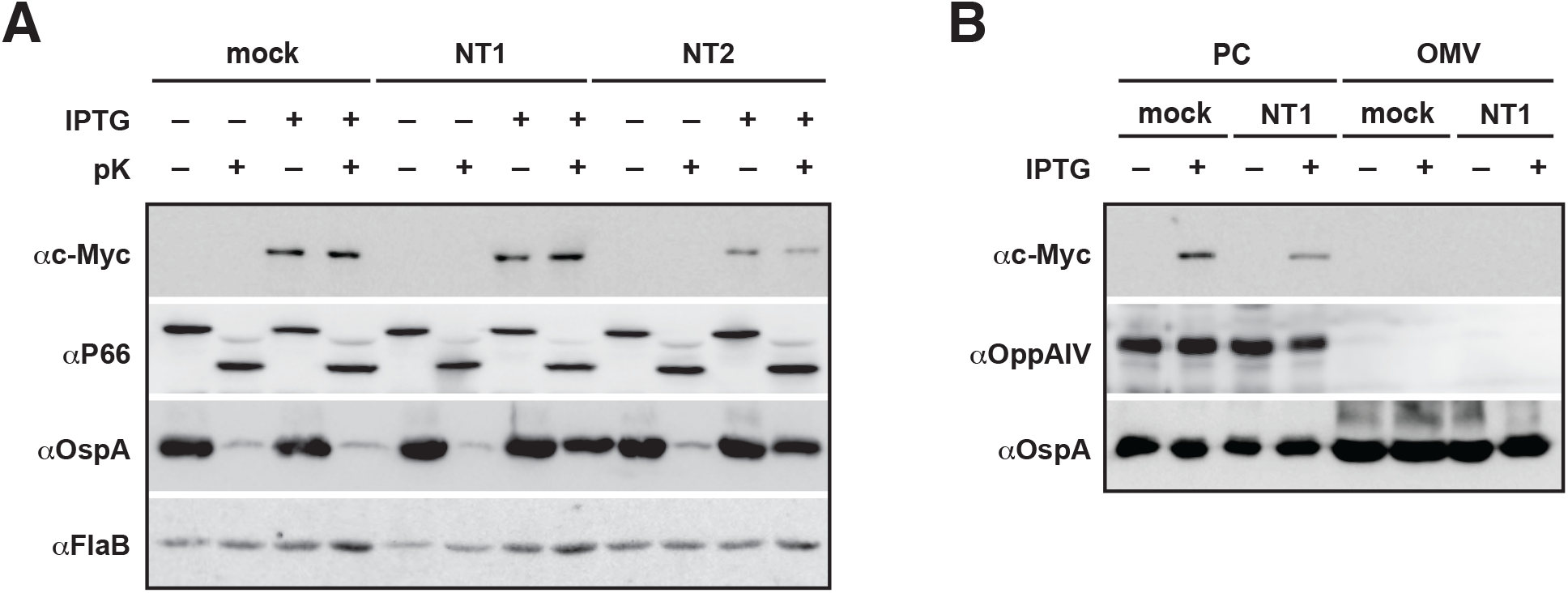
Assessment of BB0838-depended protein localization defects. **(A)** Western immunoblots of total cellular proteins of *B. burgdorferi* cells harboring “empty” pJJW101 (mock) or pJJW101 with the two BB0838-specific sgRNAs (NT1 and NT2), under non-inducing (– IPTG) or inducing (+ IPTG; 0.25mM) conditions, before (– pK) or after (+ pK) *in situ* proteolysis with proteinase K. CRISPRi induction and expression of the C-terminally c-Myc-tagged BbdCas9 was verified using an anti-c-Myc antibody. Surface lipoprotein OspA served as the model surface lipoprotein. OMP P66 served as a OMP topology control, and periplasmic FlaB was used as a cell integrity and constitutively expressed loading control. **(B)** Western immunoblots of *B. burgdorferi* outer membrane vesicle (OM) and protoplasmic cylinder (PC) fractions from *B. burgdorferi* cells harboring “empty” pJJW101 (mock) or pJJW101 with the BB0838-specific NT1 sgRNA. IM lipoprotein OppAIV and conditionally expressed dCas9-c-Myc served as OMV purity controls. Note that the PC fraction also contains OM proteins such as OspA due to the partial separation of OMVs from protoplasmic cylinders by treatment of *Borrelia* cells with hypotonic citrate buffer (Skare et al., 1995).

**Fig. 6.**
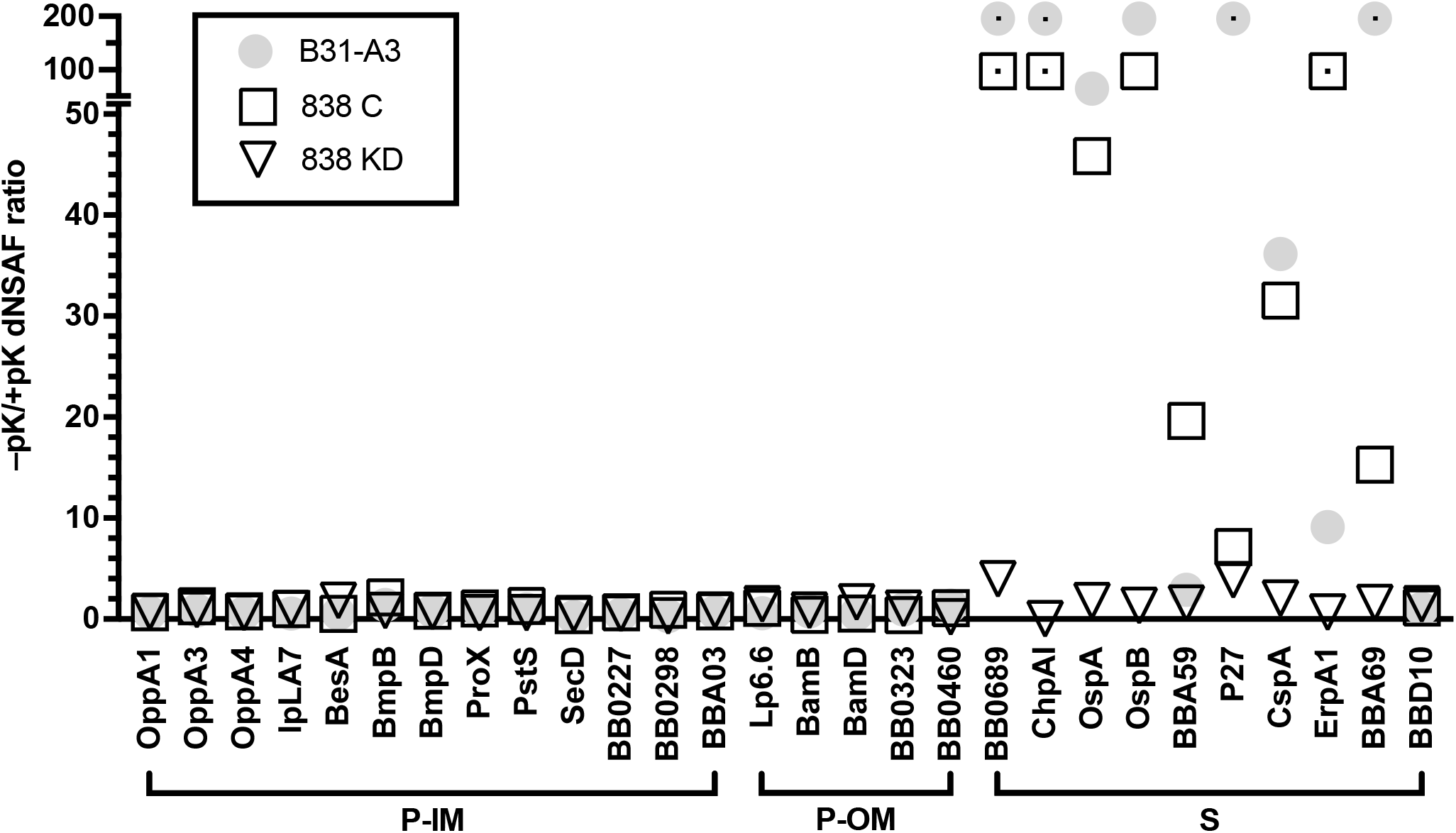
Quantitative proteomic analysis of BB0838-dependent lipoprotein localization defects. Multidimensional protein identification technology (MudPIT) quantitative proteomics analysis of membrane-associated proteins in the recombinant *B. burgdorferi* BB0838 KD strain. The plot shows the mean ratios of normalized peptide count ratios after vs. before proteinase K treatment (+pK/-pK dNSAF ratio) for subset of previously localized IM, periplasmic OM and surface lipoproteins in *B. burgdorferi* B31 clone B31-A3 (Dowdell et al. 2017). Data are shown for the inducible NT1-mediated BB0838 CRISPRi knockdown under uninduced (838C, clear squares) and induced (838KD, clear inverted triangles) conditions. Data are from 5-6 technical replicates from duplicate biological replicates. Data for B31-A3 (grey filled circles) (Dowdell et al. 2017) are shown for validation. Note that the y axis is split to help visualize the lower dNSAF ratios from low-expression proteins. See also supplementary data **Table S1 and Fig. S2**).

### Modeling of the *B. burgdorferi* Lpt pathway

In gram-negative bacteria, the LPS-transporting Lpt pathway forms a continuous periplasmic bridge between the IM and OM. A dimer of the cytoplasmic ATPase LptB forms an ABC transporter-like IM complex with integral membrane proteins LptF and LptG. This IM complex interacts via LptF with LptC, whose periplasmic domain is anchored in the IM via a single N-terminal transmembrane domain. LptC’s C-terminus itself interacts with the N-terminus of periplasmic LptA, which connects to the N-terminal periplasmic domain of LptD at the OM. LptC, LptA, and the N-terminus of LptD create a hydrophobic “greasy slide” within a continuous β -jelly-roll structure that allows for the periplasmic transport of LPS’s fatty acid membrane anchors. Having shown that BB0838/LptD_Bb_ plays a role in surface lipoprotein translocation without an apparent LptE homolog, we wondered whether *B. burgdorferi* harbors Lpt homologs upstream of LptD that would complete the pathway.

A prior search of the NCBI Protein and CDD databases by Putker et al. (Putker *et al*., 2015) using COG database categories (Tatusov *et al*., 2000) identified *B. burgdorferi* homologs for LptA (BB0465), LptB (BB0466), LptF (BB0807) and LptG (BB0808), but no clear LptC homolog. In other bacterial systems, LptC is often encoded within the same locus as LptA and LptB in an *lptCAB* operon (Putker *et al*., 2015). Based on a likely synteny in *B. burgdorferi*, we speculated that ORF BB0464 upstream of *lptA* (BB0465) could encode for the *B. burgdorferi* LptC homolog. Like LptD_Bb_, all five predicted Lpt homologs appeared to be essential based on a lack of Tn insertions (Lin *et al*., 2012, Lin *et al*., 2014).

To evaluate the five additional Lpt pathway candidates, we generated structural models using AlphaFold (Jumper & Hassabis, 2022) **(Fig. 7A)**. Signal peptides predicted by SignalP 6.0 (Teufel *et al*., 2022) were excluded. pLDDT (predicted Local Difference Distance Test) values were used as a model confidence metrics. The top model of BB0465 (LptA_Bb_) without its signal I peptide showed the expected β -jellyroll fold (pLDDT 86.4). Compared to *E. coli* LptA, BB0465 is 45 amino acids larger, which in the model extended the fold by four β -strands to 12 total. Both N and C termini appeared disordered, with potential for some short N-terminal α-helical structure. BB0464 (LptC) intriguingly was predicted to be a lipoprotein with a relatively short 14-amino acid N-terminal signal II peptide (SignalP6.0 probability = 0.99). Thus, LptC would be anchored in the IM via a different mechanism than gram-negative LptC, which uses an N-terminal α-helix membrane anchor (Wilson & Ruiz, 2022). The top model showed the expected overall β -jellyroll fold (pLDDT 90.3), with a short N-terminal α-helix following four disordered tether residues after the predicted N-terminal cysteine. The LptB (BB0466) model predictably assumed the structural fold of an ABC transporter ATPase (pLDDT 92.8). The LptF (BB0807) model showed six transmembrane helixes, with the third helix uniquely extending into the periplasm as a two-α-helix stalk before transitioning into a typical small periplasmic β -jelly roll domain (pLDDT 85.7). LptG (BB0808) modeled similarly to LptF, but lacked the periplasmic α-helical stalk (pLDDT 78.1). This analysis suggested that the *B. burgdorferi* Lpt pathway consisted of a multimeric complex of LptB_2_FGCAD, i.e., was missing only a recognizable LptE homolog found in other diderm bacterial systems.

**Fig. 7.**
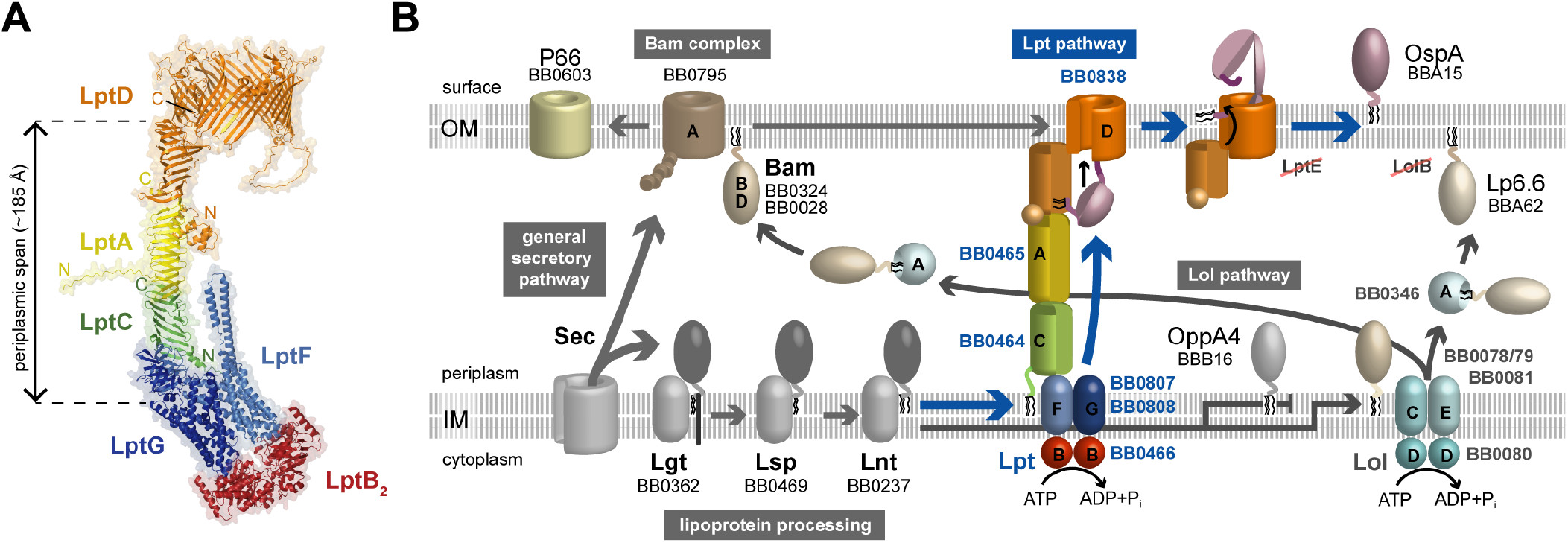
Model of *B. burgdorferi* lipoprotein transport. (A) Composite AlphaFold Multimer model of the *B. burgdorferi* Lpt pathway. Overlapping Alphafold and Alphafold Multimer models of LptD, LptD_N_A, LptD_N_AC, LptCFG and LptFGB_2_ were aligned in PyMol (see also text). Location of the subunit N and C termini are indicated in the same color as the subunits. The periplasmic span of the modeled complex was measured in PyMOL as the distance between the top of the membrane-spanning LptGF helices and the end of the periplasmic β -jelly roll domain of LptD. Note that the kink of the complex at the IM is likely an artefact of the model. **(B)** Proposed role of the *B. burgdorferi* Lpt and Lol pathways in lipoprotein sorting and secretion. *B. burgdorferi* homologs of known components are indicated by their TIGR ORF number. Canonical pathway components that are apparently missing in *Borrelia* are indicated by red strikethroughs. Model OMPs and surface, periplasmic OM and IM lipoproteins are indicated. Our model proposes that after complete posttranslational modification by the lipoprotein processing machinery in the IM, (i) *Borrelia* surface lipoproteins such as OspA are recognized and extracted by the Lpt pathway and fast-tracked along a periplasmic bridge and through the OM to the bacterial surface; (ii) the Lol pathway is solely responsible for ensuring proper localization of periplasmic OM lipoproteins such as the abundant Lp6.6 or the BAM complex associated lipoproteins BamB and BamD, and (iii) IM lipoproteins such as OppAIV avoid interaction with both the Lpt or Lol pathway to remain in the IM.

To predict protein-protein interactions and gain some initial dimensional insights, we used AlphaFold-Multimer (Evans *et al*., 2022) to generate a first model of the *B. burgdorferi* Lpt pathway. In addition to overall pLDDT, the predicted TM (pTM) and interface predicted TM (ipTM) scores were used as model confidence metrics. For multimer modeling purposes, we used only the N-terminal periplasmic domain of LptD (LptD_N_). The overlapping multimer models of LptD_N_ A (pLDDT 85.6, pTM 0.79, piTM 0.79), LptD_N_ AC (76.9, 0.70, 0.62), LptAC (78.4, 0.76, 0.71), LptCFG (76.1, 0.69, 0.64) and LptFGB_2_ (82.4, 0.78, 0.77) were assembled with the original LptD model by alignment in PyMol, resulting in the overall structural model of the *B. burgdorferi* Lpt pathway shown in **Fig. 7A**. Modeling of the entire LptB_2_ FGCAD_N_ complex resulted in the same overall structure (pLDDT 75.5, 0.8×piTM+0.2×pTM=0.545; not shown). Three key features of the predicted complex are notable: First, LptC, LptA, and the N terminus of LptD form an approximately 185 Å-long periplasmic bridge in a continual head-to-tail (N-to C-terminal) orientation. AlphaFold rejected the introduction of any additional LptA subunits into this bridge, although we were able to model LptA as a head-to-head (N-to-N terminus) homodimer and tail-to-head (C-to-N terminus) homotrimers (**Supplemental Table S3**). Second, the small α-helical domain predicted by AlphaFold at the N terminus of LptD_Bb_ moved from the monomer model orientation to now laterally interact with LptA. This suggests that this domain might function as a “clasp” to stabilize the LptA-LptD protein junction. Third, compared to LptG and other homologs, LptF contains an insertion that is predicted to protrude as an α-helical “stalk” domain into the periplasm. While this could be an artefact of the model, the DeepTMHMM transmembrane protein topology algorithm (Hallgren *et al*., 2022) equally predicted this domain to be “outside”, i.e., not part of an extended transmembrane domain.

## DISCUSSION

Envelope homeostasis is a fundamental process that ensures the continued growth and persistence of bacterial cells in a variety of sometimes hostile environments. In the arthropod-borne Lyme disease spirochete *B. burgdorferi*, this process includes the mechanisms that properly segregate over 130 lipoproteins to be either retained in the IM, transported to the periplasmic side of the OM, or secreted to the bacterial surface. Our earlier lipoproteome compartmentalization analysis showed that two-thirds, i.e. more than 87 of these lipoproteins reach the borrelial surface (Dowdell *et al*., 2017). This confirmed that secretion to the bacterial surface indeed can be considered the “default” for *B. burgdorferi* lipoproteins and suggested an efficient pathway with broad cargo specificity that connects to the posttranslational lipoprotein modification machinery in the IM and guides lipoproteins through the periplasm and the OM.

This study provided additional insights into this pathway by characterizing the biological role of *B. burgdorferi* BB0838. Both experimental evidence and structural modeling indicate that BB0838, already shown to be an integral OM protein (Kenedy *et al*., 2016), is responsible for the terminal step of surface lipoprotein secretion at the *B. burgdorferi* OM. First, the depletion of BB0838 specifically prevented the translocation of surface lipoproteins through the OM. This specific surface lipoprotein mislocalization phenotype went hand-in-hand with a significant growth defect, confirming that BB0838 is essential. Second, high-confidence *in silico* protein modeling indicated that BB0838 is a structural homolog of the gram-negative OM protein LptD, which, analogous to the secretion of amphipathic lipopolysaccharide molecules by gram-negative bacteria, would allow for the secretion of amphipathic lipoproteins in *B. burgdorferi*. The two top models consistently showed a periplasmic β -jelly roll linked to a relatively large, laterally open β -barrel pore domain that would allow for the passage of both hydrophobic and hydrophilic lipoprotein moieties and insertion into the surface leaflet of the OM lipid bilayer. However, the models differed in their integration of BB0838’s C-terminal extension that contributes to its significantly larger size (1146 amino acids, 120 kDa) compared to gram-negative LptD (784 amino acids, 87 kDa). The first model, built on a gram-negative LptDE complex by i-Tasser, inserted an LptE-like C-terminal plug domain from the outside into the β -barrel pore. The second model built by AlphaFold appeared less constrained and integrated the extension into a larger β -barrel with an estimated pore diameter of 3.7 nm. For comparison, *B. burgdorferi* outer membrane porin/adhesin P66 forms a pore with a deduced entry diameter of 1.9 nm and an internal constriction of 0.8 nm (Barcena-Uribarri *et al*., 2013). Thus, it is likely that BB0838 either assumes a structure that is a hybrid of the two models, or that the larger pore is obstructed by other means, e.g., by a yet-to-be-identified interacting protein or by transiting surface lipoproteins. Notably, both models are compatible with current topology data showing at least one protease-accessible surface loop as well as a protease-protected C terminus (Kenedy *et al*., 2016). Interestingly, LptD_Bb_ does not contain any Cys residues, indicating that unlike gram-negative LptD, it does not use disulfide bridges to coordinate conformational changes that lead to interaction with periplasmic binding partners (Ruiz *et al*., 2010, Chng *et al*., 2012).

BB0838 homologs appear conserved among pathogenic spirochetes. Based on BlastP searches, Lyme disease *Borrelia* homologs share over 90% amino acid identity, whereas relapsing fever *Borrelia* homologs (over 60% amino acid identity) and homologs found in other spirochetes such as treponemes and leptospires (around 20% amino acid identity) are more distantly related (**Table S1**). Structure homology searches revealed LptD homologs in *Treponema pallidum* (TP0515) and *Leptospira interrogans* (LIC1458). While of intermediate size, they are also predicted to fold into a similar three-domain structure (Hawley *et al*., 2021), whereas LptE homologs were only found in the LPS-displaying leptospires. It is therefore tempting to speculate that BB0838’s C-terminal domain is supporting the translocation of lipoproteins through the OM in all three spirochetal systems. At this point, we cannot exclude that spirochetal Lpt pathways transport surface glycolipids, as suggested by Hawley et al. (Hawley *et al*., 2021). However, this would clash with the rather specific phenotype of the BB0838 KD described here and the presence of glycolipids in lipid rafts of both the IM and OM of *B. burgdorferi* (Toledo *et al*., 2014, Toledo *et al*., 2015, Toledo *et al*., 2018a, Toledo *et al*., 2018b).

Multimer modeling of the predicted *B. burgdorferi* Lpt pathway complex suggests a continuous periplasmic bridge that is formed by head-to-tail monomers of LptC, LptA, and the N-terminus of LptD that is estimated to be about 18 nm long (**Fig. 7A**). This distance corresponds to the average distance between the *B. burgdorferi* IM and OM as determined by cryo-ET (Charon *et al*., 2009). Yet, periplasmic widths vary depending on the presence of flagella (Kudryashev *et al*., 2009), and the longitudinal dimensions of the periplasm-spanning *B. burgdorferi* Tol-like BesABC complex (Bunikis *et al*., 2008, Greene *et al*., 2013) and the BAM complex-associated, IM-anchored TamB (Iqbal *et al*., 2016) are modeled to be slightly larger. Thus, stoichiometric studies will have to determine whether additional LptA_Bb_ subunits, each extending the periplasmic bridge by about 6 nm, are needed to form functional multiprotein Lpt complexes.

We propose that BB0838/LptD_Bb_ acts as the previously proposed lipoprotein flippase at the end of a separate surface lipoprotein secretion system. Our data indicate that depletion of BB0838/LptD_Bb_ leads to a specific OM translocation defect of surface lipoproteins. This defect is reminiscent of the phenotypes we obtained in our previous lipoprotein localization studies, where mutations in the N-terminal disordered tether peptides of various *Borrelia* surface lipoproteins led to their mislocalization to the periplasmic leaflet of the OM (Schulze & Zückert, 2006, Schulze *et al*., 2010, Kumru *et al*., 2010, Kumru *et al*., 2011). The ability to redirect these tether mutants to the surface by destabilizing their tertiary structures (Schulze *et al*., 2010, Chen *et al*., 2011) suggested that these mutants were equivalent to secretion intermediates in the periplasmic leaflet of the OM rather than proteins that were ejected from the *Borrelia* surface lipoprotein secretion pathway at the OM. In light of the involvement of BB0838/LptD_Bb_ in surface lipoprotein secretion, we need to re-evaluate these conclusions. First, the *Borrelia* Lpt machinery with LptD_Bb_ at its terminus would provide a continuous conduit for lipoproteins to reach the surface, with limited opportunities for the egress of stalled proteins after entry at the IM. Second, surface lipoproteins interacting with the Lpt machinery would bypass the canonical OM lipoprotein sorting Lol pathway in its entirety, which would result in distinct cargo profiles for the Lpt and Lol pathways. Recent studies in *E. coli* have posited that one of the Lol pathway’s roles is to alleviate envelope stress by removing mislocalized periplasmic lipoproteins from the IM to the OM (Grabowicz & Silhavy, 2017). It is therefore possible that at least some of the lipoprotein tether mutants are rejected by the Lpt pathway at the IM and rerouted through Lol to the inner leaflet of the OM.

An earlier finding that depletion of the *B. burgdorferi* BamA ortholog BB0795 led to a decrease of both OMPs and lipoproteins in the OM (Kenedy *et al*., 2016) is compatible with the expected BAM complex dependence of BB0838/LptD_Bb_ insertion and folding. Our finding that P66, a major *B. burgdorferi* OM porin and adhesin (Bunikis & Barbour, 1999, Bunikis *et al*., 1998, Bunikis *et al*., 1995, Bunikis *et al*., 1996, Coburn & Cugini, 2003, Kenedy *et al*., 2014, Skare *et al*., 1997) maintains its w.t. topology in the absence of BB0838/LptD_Bb_ further corroborates that BB0838 acts downstream of BAM and indicates that BB0838 depletion does not lead to a more generalized OM disturbance.

The fact that BB0838/LptD_Bb_ is an essential, partially surface-exposed OMP makes it an attractive therapeutic target. LptD proteins have been successfully evaluated as potential vaccinogens for *Neisseria gonorrhoeae* and *Vibrio parahaemolyticus* (Zielke *et al*., 2016, Zha *et al*., 2016). Moreover, two novel antimicrobials targeting *Pseudomonas aeruginosa* and *E. coli* LptD have been identified. The β -hairpin-like peptidomimetic L27-11 targeting *P. aeruginosa* LptD showed potent antimicrobial activity in a mouse septicemia model (Srinivas *et al*., 2010) while the JB-95 β -hairpin macrocyclic peptide bound to *E. coli* LptD and BamA and showed potent antimicrobial activity against a panel of clinical strains (Urfer *et al*., 2016). The β-hairpin structure of these peptidomimetics might interact with the N terminus β-jellyroll of LptD (Andolina *et al*., 2018).

In summary, we used CRISPRi-mediated protein depletion to establish BB0838 as an LptD homolog that is involved in the terminal step of lipoprotein secretion to the *B. burgdorferi* surface. This suggests that the substrate specificity of Lpt pathways in diderm bacteria extends beyond LPS. Supporting this is the notion that LptD orthologs were also found in *T. pallidum* and other diderm LPS-deficient bacteria such as *Novosphingobium aromaticivorans, Deinococcus radiodurans, Thermus thermophilus*, and *Thermotoga maritima* (Putker *et al*., 2015). Yet, as in other diderm bacterial systems, *B. burgdorferi* LptD is involved in the display of lipidated immunodominant virulence factors, and thus is part of an elemental envelope homeostasis pathway that is indispensable for microbial pathogenesis. We propose that the presence of two distinct lipoprotein transport pathways in *B. burgdorferi* hints at an emerging dichotomy between Lol-mediated periplasmic lipoprotein sorting and Lpt-mediated surface lipoprotein secretion (**Fig. 7B**). This would indicate that sorting of the diverse *B. burgdorferi* lipoproteome to its three possible destinations occurs at the IM: (i) surface lipoproteins such as OspA would be fast-tracked after being recognized by LptBFG and pushed along the LptCAD periplasmic bridge and through the LptD lumen and lateral opening to the bacterial surface; (ii) periplasmic OM lipoproteins such as Lp6.6 or the essential BAM associated lipoproteins BamB and BamD would avoid interaction with the LptBFG complex to be recognized by LolCDE and transported and inserted into the periplasmic leaflet of the OM by LolA; and (iii) IM lipoproteins such as the oligopeptide-binding OppAIV would avoid either pathway to remain in the IM (**Fig. 7B**). Proper insertion and topology of LptD as the final component in the Lpt complex would be extrinsically linked to the Sec- and Lol-dependent assembly of the BAM complex in the OM. Yet, a requirement for LptC lipidation may represent an additional early and intrinsic quality control checkpoint at the IM that prevents assembly of a functional Lpt complex if its own maturation – and that of its lipoprotein cargo – is disturbed. At the same time, a lipidated LptC would lack the recently discovered rate-modulating function of the anchoring transmembrane α-helix of gram-negative LptC (Wilson & Ruiz, 2022). Our future studies will test these hypotheses and determine structure-function relationships of the *B. burgdorferi* Lpt and Lol pathway components, define their individual cargo specificities and potential interactions, and reassess previously identified *Borrelia* lipoprotein sorting determinants.

## EXPERIMENTAL PROCEDURES

### Bacterial strains and culture conditions

*E. coli* strain NEB5 (New England Biolabs) was used for plasmid construction and propagation, and grown in LB broth or on LB agar supplemented with selective antibiotic (40 µg/ml kanamycin; Sigma) as indicated. *Borrelia burgdorferi* B31-e2, a non-infectious clonal derivative of type strain B31 (Babb *et al*., 2004) encoding for 85 lipoproteins due to its reduced plasmid content (cp26, cp32-1, cp32-3, cp32-4, lp17, lp38, and lp54), was cultured in liquid or solid BSK-II medium at 34°C under a 5% CO_2_ atmosphere (Barbour, 1984, Zückert, 2007). Selective BSK-II medium contained 200 μg/ml kanamycin. Protein expression from hybrid *lac* promoters was induced by addition of isopropyl-b-D-thiogalactopyranoside (IPTG 0.25 mM; Sigma) where indicated.

### Recombinant plasmid construction and sgRNA design for CRISPRi

pJJW101 (Murphy *et al*., 2022) is a derivative of the *E. coli-B. burgdorferi* shuttle vector pJSB142 and the Mobile-CRISPRi plasmid pTn7C107 (Blevins *et al*., 2007, Peters *et al*., 2019). The plasmid encodes for an IPTG-inducible, codon-optimized non-toxic variant of SpydCas9 (BbdCas9) as well as an IPTG-inducible single guide RNA (sgRNA) cassette that allows for easy insertion of sgRNA spacer sequences. BB0838-specific sgRNA spacer sequences were designed using the web-based CRISPy-web interface (Blin *et al*., 2016) on an artificial genomic contig of *B. burgdorferi* B31 as described (Murphy *et al*., 2022). Four target sgRNAs, two complementary to the non-template (NT) strand, and two complementary to the template (T) strand were selected (**Table 2**), synthesized as complementary pairs of single-stranded oligonucleotides with BsaI site overhangs (**Table 1**), annealed, and directionally ligated into the cut *Bsa*I site array within the sgRNA cassette of pJJW101 as described (Murphy *et al*., 2022).

**Table 2.**
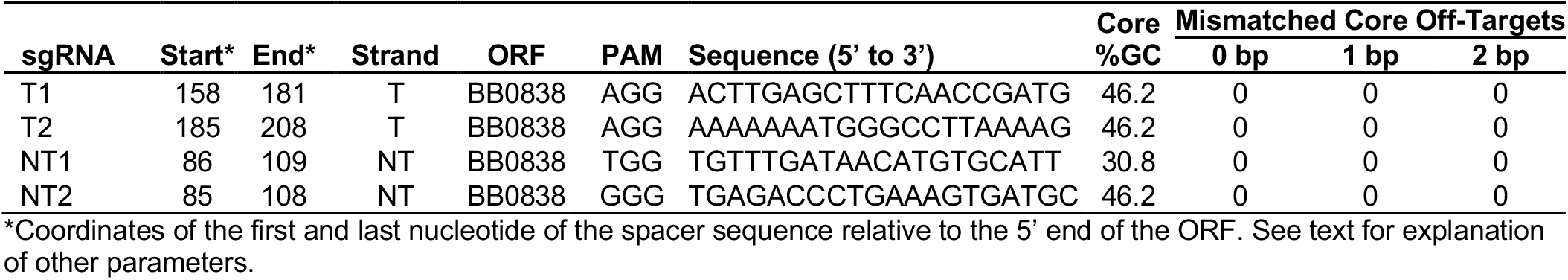
sgRNA spacer sequences used in this study.

### Growth curve assays and phase contrast microscopy

*B. burgdorferi* cells were grown from frozen stocks in selective liquid BSK-II liquid medium containing 200 µg/ml kanamycin to mid-exponential phase (about 2 days). Bacteria were then seeded at final concentrations of 1×10^5^ organisms/ml into liquid medium containing kanamycin (200 µg/ml) without or with IPTG (final concentration of 0.25 mM) and incubated at 34°C for 3 days. Spirochete numbers were determined by counting in a Petroff-Hausser counting chamber under phase contrast microscopy (Nikon Eclipse E400). For phenotypic analysis, cells were collected at day 2 post-inoculation, washed with Dulbecco’s phosphate-buffered saline (PBS, pH 7.4), fixed with 4% formaldehyde for 15 min at room temperature, and washed again with PBS to remove the formaldehyde. Micrographs were taken under phase contrast using a Nikon Eclipse E600 microscope (40× 0.55 numerical aperture Ph2 phase-contrast objective) connected to an INFINITY 3 digital camera (Teledyne Lumenera).

### Total RNA extraction and qRT-PCR analysis

*B. burgdorferi* cells grown as described above were harvested by centrifugation at 8,000 × *g* for 20 min at room temperature, washed once by resuspension in sterile room-temperature PBS containing 5 mM MgCl_2_ (PBS+Mg) and repelleted. Total RNA was extracted with TRIzol (Invitrogen) according to the manufacturer’s instructions. To remove residual DNA, the RNA samples were treated with 0.1 U DNase I (Invitrogen) for 1h at 37ºC, followed by phenol-chloroform extraction (Ambion) and ethanol precipitation overnight at -20ºC (Sambrook & Russell, 2001). The purified RNA was quantified using a Nanodrop ND1000 spectrophotometer. RNA samples were then subjected to qRT-PCR of BB0838 and *flaB* transcripts using the Luna Universal One-Step RT-qPCR kit (NEB) and an ABI Prism 7500 system (Applied Biosystems) according to the manufacturer’s instructions. Oligonucleotide primers are listed in **Table 1**. BB0838 transcript levels were validated and normalized against *flaB* mRNA, and fold changes were calculated using the comparative CT (2^-ΔΔCT^) method for quantification.

### SDS-PAGE and immunoblotting

*B. burgdorferi* cells were harvested by centrifugation and washed with PBS+Mg as described above. Bacterial pellets were solubilized in 1× sodium dodecyl sulfate-polyacrylamide gel electrophoresis (SDS-PAGE) sample buffer containing 50 mM dithiothreitol (DTT), boiled for 5 min, and stored at -20 °C. Whole cell proteins were separated on a 12% polyacrylamide SDS-PAGE gel (Sambrook & Russell, 2001) and visualized by Coomassie blue staining (Fisher BioReagents™ EZ-Run™ Protein Gel Staining Solution, #BP3620-1). For immunoblots, proteins proteins were electrophoretically transferred to nitrocellulose membranes (Millipore) using a Transblot semi-dry transfer cell (Bio-Rad). The membranes were rinsed in Tris-buffered saline (TBS) (20 mM Tris, 500 mM NaCl, pH 8.0). TBS with 0.05% Tween 20 (TBST) containing 5% dry milk was used for membrane blocking and subsequent incubation with primary and secondary antibodies; TBST alone was used for the intervening washes (Sambrook & Russell, 2001). Antibodies used were anti-OspA mouse monoclonal antibody H5332 (1:100 dilution) (Barbour *et al*., 1983), anti-c-Myc mouse monoclonal antibody (1:1000, Thermo Fisher, 9E10), anti-P66 rabbit polyclonal antibody (1:100) (Bunikis *et al*., 1995), and anti-OppAIV rabbit polyclonal antibody (1:100) (Bono *et al*., 1998). Secondary antibodies were Alkaline Phosphatase (AP)-conjugated anti-mouse (1:30,000, A3562, Sigma), and anti-rabbit antibodies (1:30,000, A3687, Sigma). Blots were developed using AP substrate CDP-Star (BioRad) for chemiluminescent detection, and signals were detected and captured using a Fujifilm LAS-4000 CCD imaging system.

### Surface proteolysis of intact *B. burgdorferi* spirochetes

Proteolytic shaving of intact spirochetes with proteinase K was performed as described (Bunikis & Barbour, 1999, Zückert *et al*., 2004, Schulze & Zückert, 2006). Briefly, *B. burgdorferi* cells harvested and washed as described above were resuspended in PBS+Mg without or with proteinase K (Invitrogen, 200 µg/ml final concentration). Proteinase K-containing and control samples were incubated for 1 h at room temperature, and reactions were stopped after 1 h by adding phenylmethylsulfonyl fluoride (PMSF) to a final concentration of 5 mM. Subsequently, cells were pelleted by centrifugation, resuspended in 1× SDS-PAGE sample buffer, boiled for 5 min, and stored at -20°C for further analysis.

### Membrane fractionations

*B. burgdorferi* outer membrane vesicles (OMVs) and protoplasmic cylinders (PCs) were isolated as described (Skare *et al*., 1995, Lenhart & Akins, 2010). Briefly, cells were harvested at room temperature by centrifugation for 20 min at 8,000 × *g*, washed in PBS with 0.1 % bovine serum albumin (BSA), and repelleted. The pellet was then resuspended in 38 ml ice-cold 25 mM citrate buffer (pH 3.2) with 0.1% BSA and incubated at room temperature for 2 hours with agitation and a 1-min vortexing step every 30 min. Next, cells were pelleted by centrifugation at 20,000 × *g* for 30 min and resuspended in 6 ml ice-cold 25 mM citrate buffer (pH 3.2) with 0.1% BSA. This sample was then layered on a discontinuous 56% (wt/wt in 4 ml), 42% (wt/wt in 15.5 ml) and 25% (wt/wt in 12.5ml) sucrose gradient in 25 mM citrate buffer in 38 ml Ultraclear tubes (Beckman-Coulter, 344058). The gradient was centrifuged at 100,000 × *g* for 18 h at 4°C (Beckman-Coulter XPN-80 Ultracentrifuge, SW32 Ti swinging-bucket rotor) to separate the OMV (upper band) and PC (lower band) fractions. The PC fraction was collected, diluted with PBS, repelleted at 20,000 × *g* for 20 min, and resuspended in 1 ml PBS with 1mM PMSF for storage at -20°C. The OMV fraction was collected, diluted with PBS, repelleted at 100,000 × *g* for 4 h at 4°C, and resuspended in 100 µl PBS for storage at -20°C.

### Quantitative mass spectrometry (MudPIT)

*B. burgdorferi* cells were subjected to surface proteolysis with proteinase K as described above and harvested. Membrane-associated proteins were then enriched by overnight extraction with Triton X-114, as described (Carroll, 2010, Dowdell *et al*., 2017). The washed detergent extracts were then precipitated at -20 ºC overnight in final 80 % (vol/vol) acetone, resuspended in 0.1 M Tris-HCl (pH 8.5), and precipitated again overnight using 20% trichloroacetic acid (TCA). The addition of acetone precipitation in the protocol was necessary to effectively remove detergent prior to analysis by MudPIT (Dowdell *et al*., 2017). Desiccated frozen protein samples from three biological replicates were then submitted for MudPIT analysis (Proteomics Center, Stowers Institute for Medical Research, Kansas City, MO). Resuspended protein samples were digested with endoproteinases Lys-C (Roche) and trypsin (Promega) at 0.1 µg/µl final concentration each. The proteinase-digested samples were then analyzed by MudPIT on an LTQ linear ion trap (Thermo Scientific) coupled to a Quaternary Agilent 1100 series high-performance liquid chromatograph (HPLC) (Florens & Washburn, 2006). Protein content in mock control versus proteinase K-treated whole-cell protein preparations was analyzed by comparison of the average distributed normalized spectral abundance factor (dNSAF) for each unique protein, which correlates directly with the relative abundance of a particular protein in the sample (Zhang *et al*., 2010). A mean dNSAF ratio of untreated control to protease-treated sample (dNSAF -pK/+pK ratio) was calculated for each protein. All MudPIT raw datasets have been deposited in the MassIVE Repository (ftp://MSV000089990@massive.ucsd.edu) and will also be available after publication from the Stowers Original Data Repository at https://www.stowers.org/research/publications/libpb-1732.

### Bioinformatics and molecular modeling

NCBI BLASTP (Johnson *et al*., 2008) was used to identify protein homologs among pathogenic spirochetes. The original structural model for BB0838 was generated by homology modeling using i-Tasser and the *Shigella flexneri* LPS-assembly protein LptD (PDB accession number 4Q35) (Qiao *et al*., 2014) as a template. Monomer models were generated using AlphaFold-monomer (Jumper & Hassabis, 2022) and assessed based on pLDDT scores and visual inspection. Protein complex models were generated using AlphaFold-Multimer (Evans *et al*., 2022) and assessed based on the weighted sum of pTM and piTM values (the AlphaFold-Multimer ranking metric), pLDDT scores, and visual inspection. Model PDB files were visualized using PyMOL (The PyMOL Molecular Graphics System, Version 2.0 Schrödinger, LLC).

## ACKNOWLEDGMENTS

We thank Sven Bergström (Umeå University, Sweden) and Patricia Rosa (NIH/NIAID Rocky Mountain Laboratories, Hamilton, Montana, USA) for antibodies and Jacob J. Wiepen for technical assistance. This work was supported in part by a KUMC Biomedical Research Training Program fellowship to HH, as well as National Institutes of Health grants P20 GM113117 (Pilot grant) and R21AI144624 to WRZ. SKS and LF were supported by the Stowers Institute for Medical Research.

## AUTHOR CONTRIBUTIONS

WRZ and HH conceived and designed the study; HH, AP, SKS and DKJ acquired the data; HH, SKS, DKJ, LF and WRZ analyzed and interpreted the data; WRZ and HH wrote the manuscript.

## GRAPHICAL ABSTRACT

Model figure based on Fig. 7

## GRAPHICAL ABSTRACT TEXT

CRISPRi was used to efficiently knock down expression of *Borrelia burgdorferi* BB0838, an essential spirochetal outer membrane protein related to LptD lipopolysaccharide (LPS) transporters in gram-negative bacteria. The BB0838 knockdown showed a specific defect in translocation of surface lipoproteins through the spirochetal outer membrane. This suggests that BB0838 functions as an outer membrane lipoprotein flippase, expanding the substrate specificity of LptD family proteins in LPS-deficient systems as part of a dichotomous Lpt/Lol lipoprotein transport system.

## PLAIN LANGUAGE SUMMARY

The surface of the Lyme disease spirochete *Borrelia burgdorferi* is covered with a variety of abundant lipoproteins, which play important roles during natural transmission and human infection following a tick bite. Here, we show that the bacterium has seemingly adopted an orphan LPS endotoxin secretion machinery found in gram-negative bacteria to secrete its complex surface lipoproteome.

## SUPPLEMENTAL DATA

*(see separate file)*

**Table S1**. BB0838 conservation among pathogenic spirochetes

**Table S2**. MudPIT data for 28 selected *B. burgdorferi* lipoproteins

**Table S3**. AlphaFold and AlphaFold Multimer models of full-length *B. burgdorferi* B31 LptD and LptA homologs

**Fig. S1**. MudPIT data for 28 selected *B. burgdorferi* lipoproteins

