## Supplemental Data (Tables S1-S3, Fig. S1) for "A *Borrelia burgdorferi* LptD Homolog Facilitates Flipping of Surface Lipoproteins Through the Spirochetal Outer Membrane"

#### Table S1. BB0838 conservation among pathogenic spirochetes

| Species and strain | Identity | NCBI Reference Sequence |
|---|---|---|
| **Lyme disease** | | |
| *Borrelia burgdorferi* B31 | 100% | WP_010889837.1 |
| *Borrelia mayonii* MN14-1420 | 94.24% | WP_075552472.1 |
| *Borrelia afzelii* ACA-1 | 91.30% | WP_004789632.1 |
| *Borrelia garinii* PBr | 91.21% | WP_004792444.1 |
| **Tick borne relapsing fever** | | |
| *Borrelia parkeri* HR1 | 65.98% | WP_038452334.1 |
| *Borrelia hermsii* DAH | 64.76% | WP_012422585.1 |
| *Borrelia turicatae* BTE5EL | 65.37% | WP_119024324.1 |
| *Borrelia hispanica* CRI | 63.80% | WP_038359268.1 |
| *Borrelia crocidurae* Achema | 63.54% | WP_014696653.1 |
| *Borrelia duttonii* Ly | 63.54% | WP_012538578.1 |
| *Borrelia miyamotoi* CA17-2241 | 61.42% | WP_117375192.1 |
| **Louse borne relapsing fever** | | |
| *Borrelia recurrentis* A1 | 63.49% | WP_012539211.1 |
| **Syphilis** | | |
| *Treponema pallidum* DAL-1 | 24.28% | WP_010881964.1 |
| **Leptospirosis** | | |
| *Leptospira weilii 56105* | 23.16% | WP_061224415.1 |
| *Leptospira borgpetersenii Brem 307* | 23.08% | WP_002730822.1 |
| *Leptospira alstonii 79601* | 22.71% | WP_020773472.1 |
| *Leptospira noguchii Bonito* | 22.51% | WP_004448668.1 |
| *Leptospira santarosai U160* | 22.51% | WP_046692483.1 |
| *Leptospira kirschneri H1* | 22.30% | WP_004765191.1 |
| *Leptospira interrogans 401* | 22.30% | WP_082271897.1 |
| *Leptospira alexanderi L60* | 22.14% | WP_039942413.1 |
| *Leptospira meyeri Veldrot Semarang 173* | 18.53% | WP_020777371.1 |

Sequence homologs and percent amino acid identities compared to *B. burgdorferi* B31 BB0838 were identified by BlastP (Johnson *et al.*, 2008).

### Table S2. MudPIT data for 28 selected *B. burgdorferi* lipoproteins *(.xlsx file)*

| | | | | dNSAF averages | | | | | | | | | | | | dNSAF ratios | | | | | |
|---|---|---|---|---|---|---|---|---|---|---|---|---|---|---|---|---|---|---|---|---|---|
| Locus | ORF | Protein Name | Localization | 838C dNSAF AVG | 838C Detected # Out of 5 | 838C_PK dNSAF AVG | 838C_PK Detected # Out of 5 | 838KD dNSAF AVG | 838KD Detected # Out of 6 | 838KD_PK dNSAF AVG | 838KD_PK Detected # Out of 5 | BbB31A3_Control dNSAF AVG | BbB31A3_Control Detected # Out of 2 | BbB31A3_ProK dNSAF AVG | BbB31A3_ProK Detected # Out of 2 | BbB31A3_Control: BbB31A3_ProK | 838C: 838C_PK | 838KD: 838KD_PK | 838KD_PK: 838C_PK | 838C: 838KD | BbB31A3_Control: 838C |
| AAC66708 | BB_0328 | **OppA1** | P-IM | 0.022405 | 5 | 0.033254 | 5 | 0.018657 | 6 | 0.025374 | 5 | 0.027067 | 2 | 0.044527 | 2 | 0.61 | 0.67 | 0.74 | 0.76 | 1.20 | 1.21 |
| AAC66706 | BB_0330 | **OppA3** | P-IM | 0.014639 | 5 | 0.011394 | 5 | 0.010294 | 6 | 0.010959 | 5 | 0.033638 | 2 | 0.038629 | 2 | 0.87 | 1.28 | 0.94 | 0.96 | 1.42 | 2.30 |
| AAC66315 | BB_B16 | **OppA4** | P-IM | 0.023723 | 5 | 0.031418 | 5 | 0.019055 | 6 | 0.029545 | 5 | 0.018812 | 2 | 0.032735 | 2 | 0.57 | 0.76 | 0.64 | 0.94 | 1.24 | 0.79 |
| AAC66748 | BB_0365 | **IpLA7** | P-IM | 0.051897 | 5 | 0.052447 | 5 | 0.055829 | 6 | 0.055025 | 5 | 0.037724 | 2 | 0.068464 | 2 | 0.55 | 0.99 | 1.01 | 1.05 | 0.93 | 0.73 |
| AAC66527 | BB_0141 | **BesA** | P-IM | 0.000281 | 4 | 0.000539 | 3 | 0.001254 | 4 | 0.000686 | 5 | 0.000472 | 2 | 0.000878 | 2 | 0.54 | 0.52 | 1.83 | 1.27 | 0.22 | 1.68 |
| AAC66758 | BB_0382 | **BmpB** | P-IM | 0.002955 | 5 | 0.001366 | 5 | 0.001741 | 6 | 0.002123 | 5 | 0.006162 | 2 | 0.004437 | 2 | 1.39 | 2.16 | 0.82 | 1.55 | **1.70** | 2.09 |
| AAB91505 | BB_0385 | **BmpD** | P-IM | 0.001233 | 5 | 0.001498 | 5 | 0.001463 | 6 | 0.001976 | 5 | 0.029778 | 2 | 0.037273 | 2 | 0.80 | 0.82 | 0.74 | 1.32 | 0.84 | **24.15** |
| AAC66525 | BB_0144 | **ProX** | P-IM | 0.009759 | 5 | 0.009214 | 5 | 0.007856 | 6 | 0.013101 | 5 | 0.019497 | 2 | 0.025897 | 2 | 0.75 | 1.06 | 0.60 | 1.42 | 1.24 | 2.00 |
| AAC66609 | BB_0215 | **PstS** | P-IM | 0.004983 | 5 | 0.003808 | 5 | 0.005365 | 6 | 0.008749 | 5 | 0.012375 | 2 | 0.013078 | 2 | 0.95 | 1.31 | 0.61 | 2.30 | 0.93 | 2.48 |
| AAC66993 | BB_0652 | **SecD** | P-IM | 0.000357 | 5 | 0.000921 | 5 | 0.000377 | 6 | 0.000738 | 5 | 0.000158 | 2 | 0.000834 | 2 | 0.19 | 0.39 | 0.51 | 0.80 | 0.95 | 0.44 |
| AAC66625 | BB_0227 | **BB0227** | P-IM | 0.002038 | 5 | 0.003083 | 5 | 0.002034 | 6 | 0.003627 | 5 | 0.002601 | 2 | 0.005516 | 2 | 0.47 | 0.66 | 0.56 | 1.18 | 1.00 | 1.28 |
| AAC66650 | BB_0298 | **BB0298** | P-IM | 0.000226 | 2 | 0.000246 | 2 | 0.000317 | 3 | 0.001101 | 4 | 0.000432 | 2 | 0.001431 | 2 | 0.30 | 0.92 | 0.29 | 4.48 | 0.71 | 1.91 |
| AAC66227 | BB_A03 | **BBA03** | P-IM | 0.013016 | 5 | 0.016956 | 5 | 0.022 | 6 | 0.025941 | 5 | 0.021405 | 2 | 0.031302 | 2 | 0.68 | 0.77 | 0.85 | 1.53 | 0.59 | 1.64 |
| AAC66241 | BB_A62 | **Lp6.6** | P-OM | 0.01302 | 5 | 0.011621 | 5 | 0.025127 | 6 | 0.016947 | 5 | 0.017272 | 2 | 0.030205 | 2 | 0.57 | 1.12 | 1.48 | 1.46 | 0.52 | 1.33 |
| AAC66428 | BB_0028 | **BamB** | P-OM | 0.005284 | 5 | 0.012558 | 5 | 0.004837 | 6 | 0.005585 | 5 | 0.008558 | 2 | 0.011199 | 2 | 0.76 | 0.42 | 0.87 | 0.44 | 1.09 | 1.62 |
| AAC66712 | BB_0324 | **BamD** | P-OM | 0.001399 | 5 | 0.002482 | 5 | 0.002372 | 5 | 0.001404 | 5 | 0.000836 | 2 | 0.001595 | 2 | 0.52 | 0.56 | 1.69 | 0.57 | 0.59 | 0.60 |
| AAC66700 | BB_0323 | **BB0323** | P-OM | 0.000205 | 4 | 0.000546 | 5 | 0.000718 | 5 | 0.00068 | 5 | 0.000288 | 1 | 0.000304 | 2 | 0.95 | 0.38 | 1.06 | 1.25 | 0.29 | 1.40 |
| AAC66836 | BB_0460 | **BB0460** | P-OM | 0.000351 | 3 | 0.000325 | 2 | 0.00027 | 3 | 0.001347 | 2 | 0.000502 | 2 | 0.000375 | 1 | 1.34 | 1.08 | 0.20 | 4.14 | 1.30 | 1.43 |
| AAC67038 | BB_0689 | **BB0689** | S | 0.00112 | 5 | 0 | 0 | 0.001795 | 6 | 0.000451 | 1 | 0.001033 | 2 | 0 | 0 | ∞ | ∞ | **3.98** | ∞ | 0.62 | 0.92 |
| AAC66230 | BB_A07 | **ChpAI** | S | 0.000224 | 2 | 0 | 0 | 0 | 0 | 0.000135 | 1 | 0.000265 | 1 | 0 | 0 | ∞ | ∞ | 0.00 | ∞ | ∞ | 1.18 |
| AAC66260 | BB_A15 | **OspA** | S | 0.253796 | 5 | 0.005538 | 5 | 0.303055 | 6 | 0.167777 | 5 | 0.213833 | 2 | 0.003307 | 2 | 64.66 | 45.83 | 1.81 | 30.30 | 0.84 | 0.84 |
| AAC66243 | BB_A16 | **OspB** | S | 0.182371 | 5 | 0.001868 | 5 | 0.158838 | 6 | 0.120432 | 5 | 0.057216 | 2 | 0.000293 | 2 | **195.28** | **97.63** | 1.32 | **64.47** | 1.15 | 0.31 |
| AAC66238 | BB_A59 | **BBA59** | S | 0.010297 | 5 | 0.000525 | 2 | 0.007756 | 6 | 0.005033 | 5 | 0.011252 | 2 | 0.003863 | 2 | 2.91 | 19.61 | 1.54 | 9.59 | 1.33 | 1.09 |
| AAC66239 | BB_A60 | **P27** | S | 0.001018 | 5 | 0.000143 | 1 | 0.001817 | 6 | 0.000469 | 4 | 0.002435 | 2 | 0 | 0 | ∞ | 7.12 | 3.87 | 3.28 | 0.56 | 2.39 |
| AAC66286 | BB_A68 | **CspA** | S | 0.003721 | 5 | 0.000118 | 3 | 0.006036 | 6 | 0.002947 | 5 | 0.005855 | 2 | 0.000162 | 2 | 36.14 | 31.53 | 2.05 | 24.97 | 0.62 | 1.57 |
| AAF07400 | BB_P38 | **ErpA1** | S | 0.000527 | 5 | 0 | 0 | 0.0012 | 6 | 0.001811 | 5 | 0.009091 | 2 | 0.000998 | 2 | 9.11 | ∞ | 0.66 | ∞ | 0.44 | **17.25** |
| AAC66287 | BB_A69 | **BBA69** | S | 0.002659 | 5 | 0.000174 | 2 | 0.003813 | 6 | 0.002343 | 5 | 0.003242 | 2 | 0 | 0 | ∞ | 15.28 | 1.63 | 13.47 | 0.70 | 1.22 |
| AAC66352 | BB_D10 | **BBD10** | S | 0.000879 | 5 | 0.000781 | 4 | 0.001273 | 6 | 0.000853 | 5 | 0.000479 | 2 | 0.000465 | 2 | 1.03 | 1.13 | 1.49 | 1.09 | 0.69 | 0.54 |
| | | | | | | | | | | | | Data from Dowdell et al., 2017 | | | | B31A3 | 838C | 838KD | 838KD/C | 838C/KD | B31A3/838C |
| | | | | | | | | | | | | | | | | surface exposure | | | Δ surface exp. | Δ expression levels | |

Lipoproteins that were detected at least twice in the mock control sample (838C; uninduced BB0838 KD, -pK) were selected from the full MudPIT dataset for further analysis. P-IM, periplasmic inner membrane; P-OM, periplasmic outer membrane; S, surface. Data for *B. burgdorferi* clone B31-A3 (Dowdell et al., 2017) are included for comparison. Data in the fat box were used for **Fig. 6**.

**Table S3. AlphaFold models of full-length *B. burgdorferi* B31 LptD and LptA homologs**

| Protein | Comments | AlphaFold Model |
|---|---|---|
| **LptD$_{Bb}$** | • Average pLDDT: 62.2<br>• N-terminal residues 1-31 disordered (including predicted signal I peptide 1-29)<br>• Residues 32-98 fold into a separate $\alpha$-helical domain followed by 7 disordered residues<br>• Disordered regions where barrel closes<br>• Disordered regions on long linkers between strands on the opposite side | 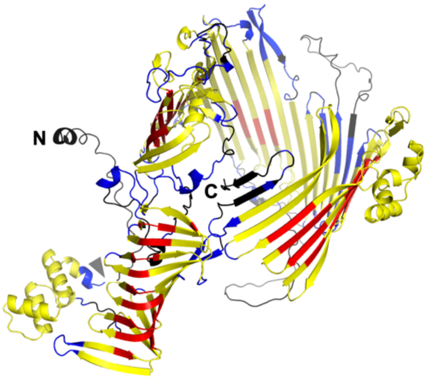  |
| **LptA$_{Bb}$** monomer | • Average pLDDT: 86.5<br>• roughly 50% longer than *E. coli* LptA (12 vs. 8 $\beta$-strands)<br>• N-terminal helix (residues 2-23, including predicted signal I peptide 1-19)<br>• Disordered N-terminal residues 24-38<br>• C terminal disordered residues | 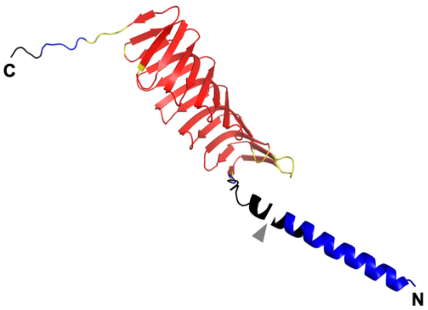  |
| **LptA$_{Bb}$** dimer | • Average pLDDT: 82.9<br>• 0.8 $\times$ ipTM + 0.2 $\times$ pTM: 0.715<br>• Stacks head-to-head (N- to N-terminus)<br>• N-terminal low confidence helix (residues 14-30, including end of signal I peptide)<br>• High confidence interface | 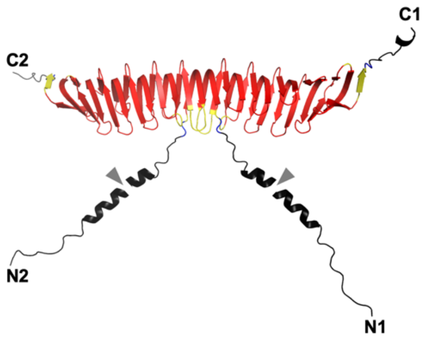  |
| **LptA$_{Bb}$** trimer | • Average pLDDT: 74.7<br>• 0.8 $\times$ ipTM + 0.2 $\times$ pTM: 0.357<br>• Stacks head-to-tail (N- to C-terminus)<br>• N-terminal low confidence short helix (residues 24-28)<br>• Lower confidence at interface | 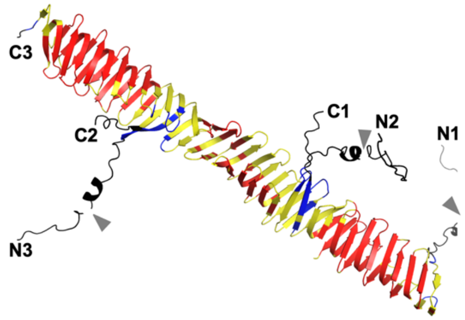  |

pLDDT (predicted Local Difference Distance Test): >90 (red): high confidence; 70-90 (yellow): confidence, 50-70 (blue): low confidence; <50 (black): disordered/no confidence. N and C termini of the LptA multimers are labeled by N# and C#, with # indicating the Lpt subunit. Gray arrows in the model figures indicate the predicted signal I peptidase cleavage sites.

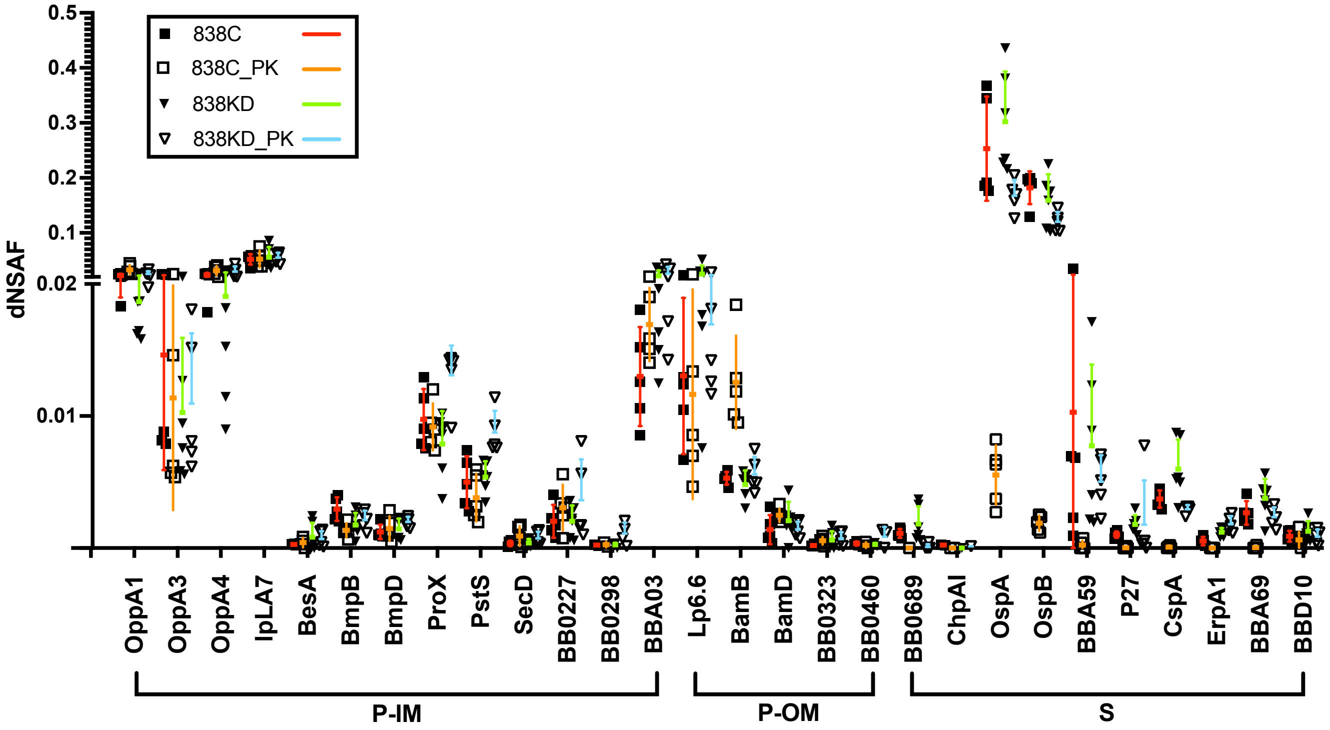


**Fig. S1. MudPIT data for 28 selected *B. burgdorferi* lipoproteins.** Plot of individual dNSAF values obtained for the 28 selected lipoproteins in 5 or 6 technical replicates. 838C, uninduced BB0838 KD, -pK; 838C_PK, uninduced BB0838 KD, +pK; 838KD, induced BB0838 KD, -pK; 838KD_PK, induced BB0838 KD, +pK). Color-coded error bars show the mean ± SD. Note that the y axis is split to help visualize the lower dNSAF ratios from lower abundance proteins. The derived dNSAF means for each protein were used in **Table S1** and **Fig. 6**.
